# Generating Agent-Based Multiscale Multicellular Spatiotemporal Models from Ordinary Differential Equations of Biological Systems, with Applications in Viral Infection

**DOI:** 10.1101/2021.01.28.428647

**Authors:** T.J. Sego, Josua O. Aponte-Serrano, Juliano F. Gianlupi, James A. Glazier

## Abstract

The biophysics of an organism span scales from subcellular to organismal and include spatial processes like diffusion of molecules, cell migration, and flow of intravenous fluids. Mathematical biology seeks to explain biophysical processes in mathematical terms at, and across, all relevant spatial and temporal scales. While non-spatial, ordinary differential equation (ODE) models are often used and readily calibrated to experimental data, they do not explicitly represent spatial and stochastic features of a biological system, limiting their insights and applications. Spatial models describe biological systems with spatial information but are mathematically complex and computationally expensive, which limits the ability to calibrate and deploy them. In this work we develop a formal method for deriving cell-based, spatial, multicellular models from ODE models of population dynamics in biological systems, and vice-versa. We provide examples of generating spatiotemporal, multicellular models from ODE models of viral infection and immune response. In these models the determinants of agreement of spatial and non-spatial models are the degree of spatial heterogeneity in viral production and rates of extracellular viral diffusion and decay. We show how ODE model parameters can implicitly represent spatial parameters, and cell-based spatial models can generate uncertain predictions through sensitivity to stochastic cellular events, which is not a feature of ODE models. Using our method, we can test ODE models in a multicellular, spatial context and translate information to and from non-spatial and spatial models, which help to employ spatiotemporal multicellular models using calibrated ODE model parameters, investigate objects and processes implicitly represented by ODE model terms and parameters, and improve the reproducibility of spatial, stochastic models. We hope to employ our method to generate new ODE model terms from spatiotemporal, multicellular models, recast popular ODE models on a cellular basis, and generate better models for critical applications where spatial and stochastic features affect outcomes.

**Statement of Significance:** Ordinary differential equations (ODEs) are widely used to model and efficiently simulate multicellular systems without explicit spatial information, while spatial models permit explicit spatiotemporal modeling but are mathematically complicated and computationally expensive. In this work we develop a method to generate stochastic, agent-based, multiscale models of multicellular systems with spatial resolution at the cellular level according to non-spatial ODE models. We demonstrate how to directly translate model terms and parameters between ODE and spatial models and apply non-spatial model terms to boundary conditions using examples of viral infection modeling, and show how spatial models can interrogate implicitly represented biophysical mechanisms in non-spatial models. We discuss strategies for co-developing spatial and non-spatial models and reconciling disagreements between them.

## Introduction

The function and form of biological systems involve biophysical mechanisms across all scales, from the subcellular to the organismal (1). Processes and properties observed at a particular scale emerge from, and affect, complex processes at a finer scale, such as the emergent polarization and translocation of a cell from complex subcellular reaction kinetics when exposed to a chemotactic stimulus (2). Such hierarchical organization can be established to relate the roles of individual biochemical components in signaling and cytoskeletal networks at the subcellular level to the emergence of tissue-level force generation, cell migration and tissue function and shape changes (3). Between these two scales lies the cell, which has been argued to provide a natural level of abstraction for modeling development (4). The role played by individual cells and their interactions is apparent in development and disease, including tip cells in collective cell migration (5), diversity in production of virions during viral infection (6) and cancer initiation by stem cells (7).

Each cell occupies a volume in space, the biochemical processes in which are well known to experience effects of spatial mechanisms like volume exclusion of macromolecules, the cytoskeleton and organelles, resulting in macromolecular crowding that causes highly non-mass-action reaction rates (8) and anomalous diffusion (9). Recent computational models have sought to describe the spatiotemporal effects on the kinetics of elementary reactions (10), as well as to relate discrete and continuous mathematical descriptions of spatially resolved subcellular reaction kinetics (11). Likewise, models of spatiotemporal multicellular systems in development and disease have shown the significance of spatial mechanisms like diffusive transport and the shape and position of individual cells in angiogenesis (12), polycystic kidney disease (13) and spheroid fusion (14). Simulations of viral infection have demonstrated the non-negligible effects of the well-mixed assumptions commonly employed when modeling viral infection and immune response using population dynamics, like the neglect of the initial distribution of infected cells in susceptible tissue (15).

The ability to derive cell-based, spatiotemporal models from ODE models would enhance the utility of both types of models. Cell-based, spatiotemporal models can explicitly describe cellular and spatial mechanisms neglected by ODE models that affect the emergent dynamics and properties of multicellular systems, such as the influence of dynamic aggregate shape on diffusion-limited growth dynamics (16) and individual infected cells on the progression of viral infection (17). Likewise ODE models can inform cell-based, spatiotemporal models with efficient parameter fitting to experimental data, and can appropriately describe dynamics at coarser scales and distant locales with respect to a particular multicellular domain of interest (*e.g.*, the population dynamics of a lymph node when explicitly modeling local viral infection). One such example is the approach of Murray and Goyal to derive discrete stochastic dynamics from continuous dynamical descriptions using the Poisson distribution in their multiscale modeling work on hepatitis B virus infection (18). However, to our knowledge, no well-defined general formalism describes systematic translation of models to the cellular scale from coarser scales at which spatially homogeneous, population dynamics models using ordinary differential equations (ODEs) appropriately describe a biological system. In the very least, the lack of consistent translation of model terms and parameters between spatial and non-spatial models severely inhibits the potential to apply the vast amount of available information and resources in non-spatial modeling, such as those available in BioModels (19), to spatial contexts, and to share validated, reproducible spatial models (20).

In this work we develop a method for generating spatial, cell-based models from homogeneous, ODE-based models of biological systems to address critically important questions such as

1. Is the behavior of an ensemble of *N* cells with mean parameter *x* in mechanism *y* the same as *N* times the behavior of a single cell with the same mechanism and mean parameter?
2. Is the behavior of a biological system in which a species realistically diffuses qualitatively different from one in which the species diffuses infinitely fast?

As such, we develop our method under the premise that a spatial, cell-based model applied to an ensemble of cells reproduces the ODE model from which it was generated in the limit of well-mixed conditions. We describe the general formalism of our method with regard to continuous modeling of cell populations and soluble signals, and spatiotemporal, multicellular and cell-based mechanisms (*e.g.*, diffusion, contact-mediated interactions, discrete cell type transition probabilities). We apply our method to generate spatial, cell-based models from two non-spatial models of viral infection and host-pathogen interaction based on our recent work in viral infection and immune response modeling (17) to show critically important aspects of spatially homogeneous vs. heterogeneous models. We call our method *cellularization*, which is to mean that cellularizing a homogeneous model generates an explicit representation of cellular aspects implicitly represented by the homogeneous model.

## Methods

In this section we describe our method for converting model objects and processes between non-spatial ODE models and spatial, cell-based models of multicellular systems. We consider paired models of the same underlying biological/biochemical system, where an ODE model consists of only scalar-valued variables related by rate laws that depend only on the variables and a set of scalar parameters, and hence is intrinsically non-spatial. The spatial model could include scalar variables, continuously variable spatial quantities (*i.e.*, fields) and discrete agents with individual states, which typically have spatial locations (which could change in type) and which can come into existence (*e.g.*, birth) and disappear (*e.g.*, death). The relationships among agents and variables are more diverse in spatial models, including the rate equations of the non-spatial case, as well as stochastic transition rules, partial-differential equations describing the effect of spatial variation on rates, and more complex boundary and initial condition considerations. Initial conditions require consist specification of initial values of variables only among paired models, with a limited number of possible boundary conditions in the spatial model. Rate laws with explicit time delays and integro-differential forms are extensions of the basic forms treated in this work, though many of the issues we consider apply. Likewise, nonspatial agent-based models form an intermediate class, which we do not consider in detail here, though many of the same considerations apply.

We define a spatial model as analogous to a non-spatial ODE model if the ODE model describes the dynamics of the spatial model in the limit of well-mixed conditions. As such, we refer to a spatial model generated from an ODE model as the spatial analogue of the ODE model. In this work we consider cells, diffusive species, and the interactions between them. We assume that both an ODE model and its analogous spatial model describe the same underlying real-world dynamics of a biological system, but that only the spatial model explicitly describes spatial dynamics. We assume that the ODE model assumed well-mixed conditions such that all ODE model variables, parameters and equations represent spatially homogeneous properties of, and interactions between, underlying objects. Likewise, we assume that the spatial domain in which the objects and processes are described by an analogous spatial model represents a constituent spatial element of the biological system described by the ODE model. Under these assumptions, the following outcomes are possible when comparing results from a pair of ODE and spatial models,

- **Homogeneity**. The models agree for the same parameters and the system is spatially homogeneous.
- **Ensemble average**. The models agree for the same parameters and the system is spatially heterogeneous.
- **Localization**. The models agree for different parameters and the system is spatially heterogeneous.
- **Incompatibilit**y. The models never agree.

While in general nothing prohibits us from defining multiple spatial domains that represent various interesting elements of the biological system, we assume the simplest case in this work when deriving analogous mathematical forms – that a spatial domain is a representative volume element of the entire biological system described by an ODE model. We describe our formalism as beginning with an existing ODE model and deriving an analogous spatial model. However, our formalism works in the opposite direction, allowing us to derive an ODE model from an existing spatial model.

### Spatiotemporal Scaling Using Well-Mixed Conditions

Well-mixed conditions allow us to relate measurements of quantities at different scales, where measurements at every scale everywhere exactly and uniformly scale. For a measurement *Z* of a quantity at one scale and *z* measuring the quantity of the same object but at another scale, let *μ* be a coefficient relating the two scales such that

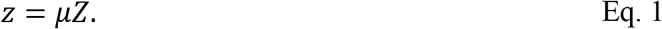

The same is true for the behavior of objects according to well-mixed conditions, where changing the scale at which an object operates does not qualitatively affect the dynamics of the object described by the ODE model. As such, for a rate equation of the form,

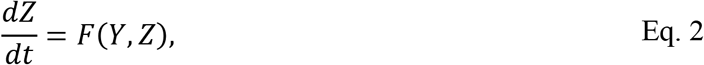

dynamics for a measurement *z* at some other scale and *y* = *μY* take the form,

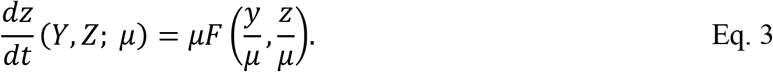

Scaling of ODE model quantities under well-mixed conditions is assumed to be true for quantities of mass and number (*e.g.*, the population of cells of a particular type) at every scale. Depending on the scale of a multiscale spatial model, well-mixed conditions allow us to cast a quantity *Z* from an ODE model as acting globally (*i.e.*, as spatially homogeneous) or locally (*i.e.*, as spatially heterogeneous) in a spatial model, depending on the spatial qualities imposed upon *Z*. For example, diffusive species with very high or very low diffusivity may be cast as acting globally or locally, respectively, at the scale of individual cells. Henceforth, for a measurement *Z* of the quantity of an object at the scale of an ODE model, let *z* be a global (*i.e.*, non-spatial) measurement of the quantity of the same object at the scale of a spatial model, and let 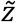 be an infinitesimal, point-like measurement of the same object.

To employ Eq. 3 to describe the dynamics of a quantity *z* at the scale of a spatial domain according to an ODE model requires the relating of a static measurement 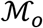 of some scalar quantity at the scale of the ODE model to a global, static measurement 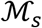 of a scalar quantity at the scale of its analogous spatial model such that, using Eq. 1, for a global scaling coefficient *η* and time *t*,

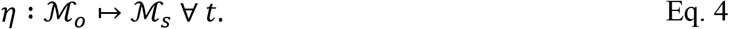

For example, an ODE model describing a volume 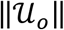 and an analogous spatial model describing a volume 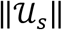 presents a form of Eq. 4 with a scaling coefficient *η*_*vol*_ = *η*,

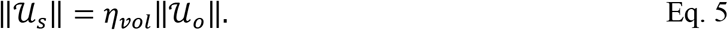

Likewise, an ODE model describing some fixed number 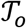 of a set of cell types and analogous spatial model describing a fixed number 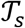 of a set of the same cell types presents a form of Eq. 4 with a scaling coefficient *η*_*count*_ = *η*,

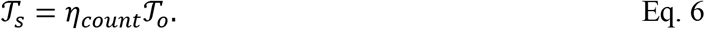

In the case of Eq. 5, application of Eq. 3 to generate global measurements at the scale of a spatial model would presuppose that, under well-mixed conditions, a quantity *Z* is the same per volume. Likewise for Eq. 6, well-mixed conditions would impose that a quantity *Z* is the same per cell.

For the case of counting numbers of a cell type in a discrete cell population, let *σ* be the identifier of a cell and let *τ*(*σ*) be the type of *σ*. Let an ODE model of the quantity of a cell type 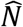 be written as *N*. Using this convention, the discrete analogue of *N* naturally follows by counting the number of cells of type 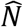,

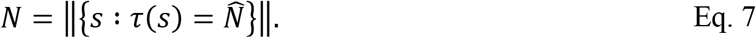

Likewise, for local measurements of some global, scalar measurement *Z* at some scale, consider a local, point-like measurement 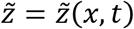 of *Z* in a space 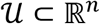, where 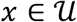. For diffusive 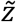, let 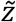 be governed by a reaction-diffusion equation of the general form,

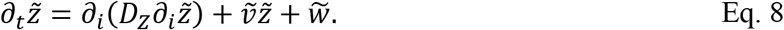

Considering the case of homogeneous 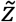 (*i.e.*, under well-mixed conditions), *Z* and 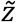 can be related with a local scaling coefficient *θ* using Eq. 1,

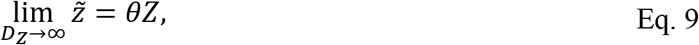

and in general, for global quantity *z* at the scale of a spatial domain (*i.e.*, *z* = *ηZ*),

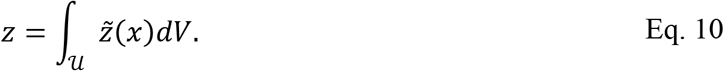

Then, when considering Eq. 10 for a homogeneous 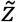,

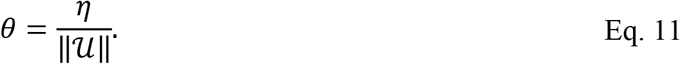

When deriving a spatial model from ODEs, the following relationships are assumed to hold under well-mixed conditions for ODE model measurement *Z*, spatial global measurement *z*, and local, point-like measurement 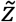, according to Eq. 5 and Eq. 11

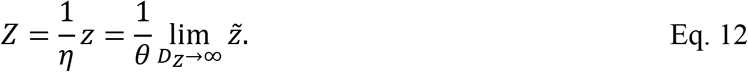

Using Eq. 12, quantities described by an ODE model at some scale can then be employed in a spatial model at some smaller scale as either globally acting objects (according to the ODE model, using Eq. 3) or a locally acting object (according to Eq. 8 for diffusive species, and to subsequent derivations for cells).

To define a point-like measurement of a quantity associated with a particular cell, well-mixed conditions are also applied to the domain of a cell. Let *σ*(*x*, *t*) be a cell identifier that describes the location of individual cells in the spatial domain (*e.g.*, for cell *s*, *σ*(*x*, *t*) = *s* at every point *x* occupied by cell *s* at time *t*), and let 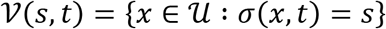 be the domain of cell *s* at time *t* in the spatial domain 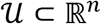. A point-like measurement 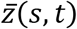 of a spatially heterogeneous quantity 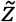 associated with a cell *s* at time *t* under well-mixed conditions is then

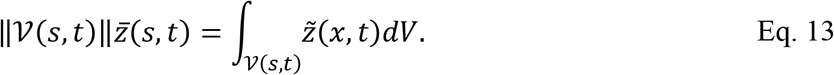

Note that for point-like cellular objects,

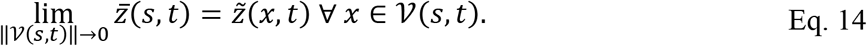

### Cellularization of Signals

Consider a rate equation for chemical species *F* and *G* and number *N* of a cell type 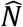,

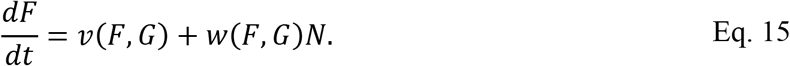

Here *v* and *w* are generic functions with arguments *F* and *G* for demonstrative purposes, but are otherwise arbitrarily selected (*i.e.*, they could be any set of chemical species or some other object of the system). Using Eq. 12, let 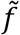 be the heterogeneous analogue of *F* according to Eq. 8 with diffusivity *D*_*F*_. In general, *v* can be applied in any spatially particular way (*e.g.*, only as a flux on a boundary) on the condition that the volume integral of its spatial analogue is equal to *ηv* at every time *t*. We consider here the case when *v* is uniformly applied in the spatial domain,

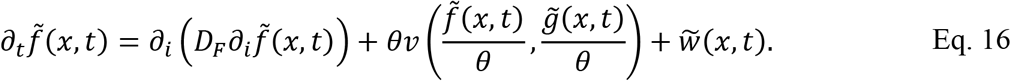

Here *v* is rewritten as a substitution of 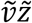 in Eq. 8 using Eq. 3. In the case of heterogeneous 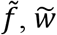 can be related to *wN* in Eq. 15 by integrating over all cell volumes of type 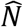,

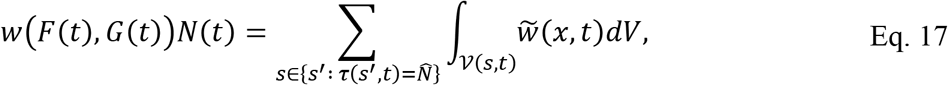

We can write an expression for 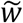 at each point as a function of 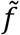 and 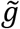 such that Eq. 17 is true under well-mixed conditions by employing Eq. 13,

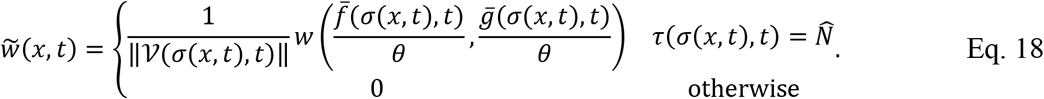

Note that *w* in Eq. 13 and 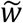 in Eq. 18 are general in the sense that multiple terms could be defined with respect to multiple cell types. In such a case, each type-specific term would have a corresponding 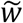 according to the general form in Eq. 17.

### Hybridization of Population Dynamics

We derive discrete, stochastic events from ODE rate equations by considering the ODE rate equations as mean-field approximations of discrete, stochastic events. The total number of occurrences of a discrete, stochastic event *K* is assumed to be a random variable drawn from a Poisson distribution, which allows us to relate expressions from ODE and discrete, stochastic models. We use the cumulative distribution function 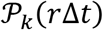 for determining the probability that a discrete event with mean rate *r* occurs more than *k* times over a simulation step of period Δ*t*,

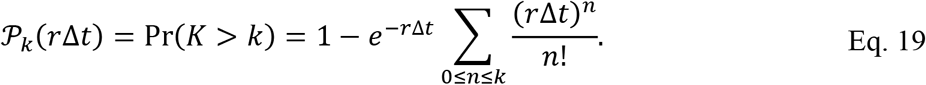

For a discrete event that can occur exactly once in a simulation step (*e.g.*, killing of one cell), we use 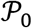, which determines the probability of an event having occurred at least once (rather than exactly once).

Consider ODE-model population dynamics of the number of cell types 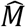 and 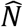 of the form,

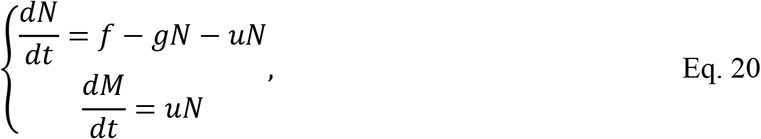

Three events are described in the rate equation for *N*. First, the term *f* describes the net inflow of 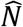 − type cells. Second, the term −*gN* describes the net outflow of 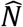 − type cells. Third, the term −*uN* describes the transition of a cell from a 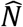 − type cell to a 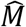 − type cell (*e.g.*, from a living cell to a dead cell), which we denote as 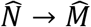. Discrete, stochastic analogues are derived in this order using a mean-field approximation of the Poisson distribution.

The net inflow of 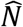 − type cells is written using the cumulative distribution function of the Poisson distribution for the occurrence of *k* cells of type 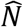 being added to the spatial domain,

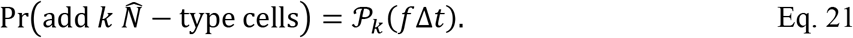

Eq. 21 is implemented using the following algorithm. Beginning with *k* = 0, draw a uniformly distributed random number *X* in [0, 1]. If *X* is greater than Eq. 21 for *k*, then add *k* cells of type 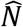. Otherwise, increment *k* by one and repeat. Functionally, this imposes that, while counting upwards from *k* = 0, if no more than *k* cells are added, then *k* cells are added.

The net outflow of 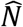 − type cells is considered for each 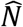 − type cell independently at each simulation step as an event that can occur no more than once. For cell *s*,

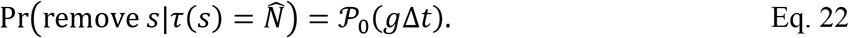

Note that a form like Eq. 22 can also be generated for describing the probability of mitosis in the case where the term −*gN* in Eq. 20 were instead +*gN*.

The discrete transition event of a cell of type 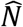 to type 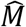 over a period Δ*t* is considered for each 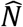 − type cell of identification *s* and type *τ*(*s*, *t*) at time *t* as an event that can occur no more than once,

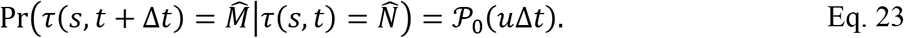

### Hybridization of Contact-Mediated Population Interactions

Contact-mediated interactions are critical to many developmental (*e.g.*, stem cell proliferation) and physiological (*e.g.*, antigen presentation) processes. In addition to the interactions described in *Hybridization of Population Dynamics*, suppose that *u* in Eq. 20 describes a rate of 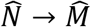 mediated by contact interactions between 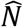 − and 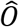 − type cells and, for *a* > 0 and cell types 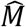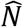, and 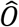, Eq. 20 has the form,

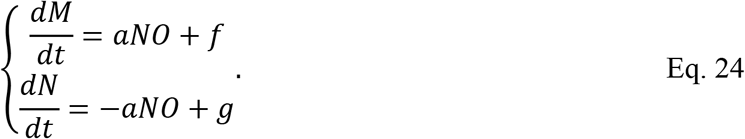

Spatial models of contact-mediated population interactions are derived by considering well-mixed conditions in the strictest sense. If 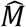 and 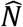 − type cells constitute a cellular body and present a certain surface area available for contact interfaces, then the surface area of contact interfaces between 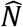 − and 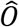 − type cells in well-mixed conditions is proportional to the number of 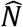 − and 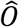 − type cells *N* and *O*, respectively.

As such, let *A*_*P*_ be the total available contact surface area of all 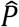 − type cells of fixed number, let *P* = *M* + *N* be a quantity of a set of 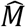 and 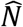 cells, let *A*_*N*_ be the total available contact surface area of a 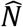 − type cell, and let *A*_*N*,*O*_ be the contact area of a 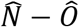 interface. Under well-mixed conditions,

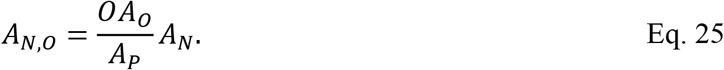

Eq. 25 can be rewritten for *O*, in which case Eq. 24 can be rewritten on a cellular basis for cell *s* with total interfacial contact area with 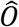 − type cells *A*_*s*,*O*_ (*i.e.*, *A*_*N*,*O*_ → *A*_*s*,*O*_ on a cellular basis). 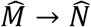 mediated by 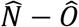 contact interfaces is then considered for each cell of identification *s* and type 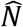 as an event that can occur no more than once,

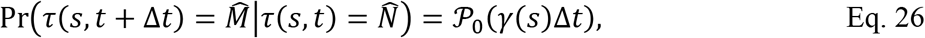

where

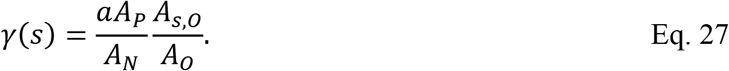

Eq. 27 scales *a* from the general ODE form in Eq. 24 to the ratio of two ratios, namely the ratio of the contact area of a cell with 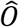 − type cells to the total contact area of a 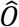 − type cell, to the ratio of the total contact area of a 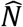 − type cell to the total contact area of all 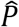 − type cells. Note that in the case of geometrically identical 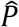 − type cells (*i.e.*, 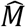 − and 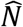 − type cells have the same total contact surface area), Eq. 27 can be rewritten to scale *a* to the total number of 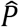 − type cells,

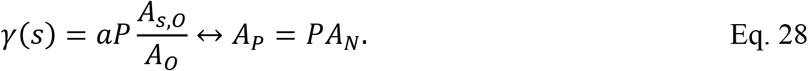

### Hybridization of Recruitment

Eq. 20 describes the inflow of cells with a rate *f* and outflow of cells with a rate *g* per cell. Here we describe handling inflow of cells, which can be generally described, and neglect describing any general method for outflow, which is particular to problem setup and cellular dynamics method (*e.g.*, whether cells are simply removed wherever they are when outflow occurs, or they first migrate towards a boundary). To describe the inflow of cells in a spatial model, it is necessary to define a boundary through which each incoming cell enters the spatial domain, and the location on that boundary where each incoming cell appears. For an inflow rate *f* into a spatial domain 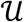, we refer to a corresponding boundary 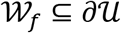 on which cells appear as an *inflow boundary*. Likewise, we refer to placing a new cell on an inflow boundary as *seeding*. A cell can be seeded at a site on an inflow boundary that is not occupied by another cell. We refer to such sites as being *available* and denote the set of available sites on an inflow boundary for an inflow rate *f* as 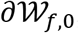,

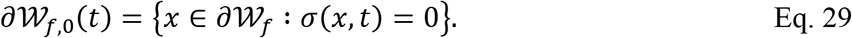

In general, we define an available site selection function 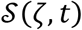 that selects an available site for seeding a cell of type *ζ* at time *t*. It should be noted that an available site selection function does not describe the probability of seeding a cell (*e.g.*, Eq. 21), but rather maps the set of available sites on inflow boundaries to a coordinate where seeding a cell occurs for a given cell type and time. It should also be noted that an available site selection function is defined to take the argument of a cell type, rather than an inflow rate, since one could define a total inflow rate as consisting of multiple inflow rates on multiple inflow boundaries (*e.g.*, two thirds of incoming cells enter on one boundary, and one third of incoming cells enter on another boundary). Imposing no additional spatial information on seeding an incoming cell results from designating all boundaries of the spatial domain as inflow boundaries, and then randomly selecting an available site with a uniform distribution. Likewise designating a subset of the boundaries of a spatial domain as inflow boundaries necessarily imposes additional spatial information onto the spatial analogue of an ODE model by introducing spatial heterogeneity to the boundaries of the spatial domain.

An available site selection function of particular interest (though not critical to the total method presented here) that we propose concerns the influence of local heterogeneity of signals in a spatial domain that induce directional migration (*e.g.*, chemotaxis, haptotaxis) in incoming cells. Suppose that a locally heterogeneous field *c*(*x*, *t*) attracts an incoming cell of type *ζ* that is to be seeded somewhere in a set of available sites 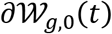 on an inflow boundary according to a rate *g*. Rather than impose additional spatial information outside of a spatial domain, we can instead presuppose that *c* near 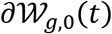 influences an otherwise random seeding location where an incoming cell of type *ζ* arrives. In this case, consider a randomly selected subset of available sites 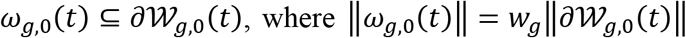 for a seeding fraction *wg* ∈ (0, 1] and each selected site *y* ∈ *ω*_*g*,0_(*t*) is selected with a uniform probability (*i.e.*, without imposing any additional spatial information, as mentioned in the preceding text). We impose the effects of a locally heterogeneous attractant *c* of cell type *ζ* at time *t* by seeding an incoming cell of type *ζ* at the location where *c* is greatest among the randomly selected subset of available sites,

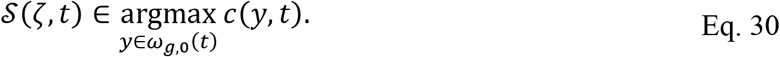

It naturally follows that replacing argmax(∙) with argmin(∙) imposes a repellant *c*. Note that additional selections must be made when this particular available site selection function yields multiple values (*e.g.*, we randomly choose a site when *c* is everywhere zero). Also note that *w*_*g*_ = 1 imposes that an incoming cell is seeded exactly at the location with the maximum value of *c* in all available sites 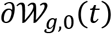, while *w*_*g*_ → 0 imposes that an incoming cell is seeded at a random location.

### Mixing Mixed Conditions

Nothing prohibits us from supposing that only some of the cells of a population act locally, while the remaining cells of the population act globally, in the sense that only the locally acting cells are represented in a spatial domain, while the effects of the entire population are present in the spatial domain. We can, for example, say that of 2*N* cells of type 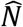, only *N* cells are present in a spatial domain, the interactions of which are modeled locally, while the other *N* cells are not present in the spatial domain, and their interactions occur everywhere homogeneously (in fact, the well-mixed conditions employed by ODE models necessitate that this is true). The same is true concerning signal fields, in that a homogeneous field can either be explicitly represented in the spatial domain, or represented as a scalar value equal to the volume integral of an equivalent homogeneous field. This distinction is important for at least two reasons.

First, from a computational standpoint, explicitly integrating the diffusion equation in time becomes increasingly computationally expensive as the diffusion coefficient (*e.g.*, *D*_*F*_ in Eq. 16) of a soluble signal increases (because of numerical stability). Likewise, as the diffusion coefficient of a soluble signal increases, the field acts more homogeneously. Other factors permitting (*e.g.*, diffusion length), one can mitigate the computational cost of integrating fast-acting soluble signals in time by representing them as scalar-valued functions (according to their ODE form) that act everywhere uniformly in a spatial domain.

Secondly, employing a cellular dynamics method that includes volume exclusion of cells introduces the possibility that no available sites exist on an inflow boundary when attempting to perform seeding. We refer to such cases as *overcrowding*. While overcrowding may elucidate problematic ODE model forms and/or parameters, we do not necessarily discard an ODE model or parameter set when overcrowding occurs (though it could inform further ODE model development), neither do we necessarily ignore a seeding event. Rather, we generally describe the inflow of cells as consisting of two stages. In the event of seeding a cell of a type *ζ*, we first add a cell to a population of *nearby* cells of type *ζ*. Nearby cells are not spatially represented, but instead act everywhere homogeneously (*i.e.*, as if they were global) in the spatial domain according to ODE model forms. Before performing all spatial interactions of a simulation step, we attempt to seed each nearby cell. Each nearby cell that is successfully seeded is then removed from the population of nearby cells, and the remaining population of nearby cells act homogeneously in the spatial domain for the simulation step.

### Implementation Details

The remaining aspects of cellularizing an ODE model depend on the choice in cellular dynamics method to explicitly describe the spatial dynamics that are implicitly represented in an ODE model. Such a choice largely depends on the scale and resolution of the spatial domain, whether on the order of microns, centimeters or otherwise, which dictates the appropriateness of a particular cellular dynamics method. Various cellular dynamics methods have their own mathematical and computational details that affect the behavior of model cells and their emergent behaviors and properties (21).

To demonstrate cellularization of ODE models in this work, we employed the Cellular Potts model (CPM, or Glazier-Graner-Hogeweg model) to model cellular dynamics in analogous spatial models. The CPM represents individual cells and a general medium as deformable, volume-excluding spatial objects in a lattice that defines a spatial domain (22), and is implemented in multicellular modeling and simulation software like CompuCell3D (23), Morpheus (24) and Chaste (25). Cell motility in the CPM is modeled as the stochastic exchanging of lattice sites at intercellular and cell-medium interfaces. In the CPM, a pair of neighboring lattice sites 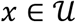 and 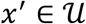 are randomly selected and the case is considered where the identification at *x*^′^ attempts to copy itself to *x*, called a copy attempt and denoted *σ*(*x*, *t*) → *σ*(*x*^′^, *t*). A copy attempt occurs with a probability according to a system effective energy 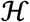 that models various cellular properties and behaviors (*e.g.*, volume constraints, adhesion, chemotaxis). For each copy attempt, the change in system effective energy 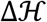 due to *σ*(*x*, *t*) → *σ*(*x*^′^, *t*) is calculated. Copy attempts that decrease the system effective energy are always accepted, and copy attempts that increase the system effective energy occur according to a Boltzmann acceptance function of the change in system effective energy,

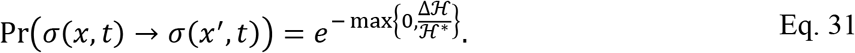

Here 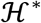 is the intrinsic random motility that affects the stochasticity of copy attempts.

In this work we employed a system effective energy modeling a volume constraint for each cell, adhesion at all interfaces and chemotaxis,

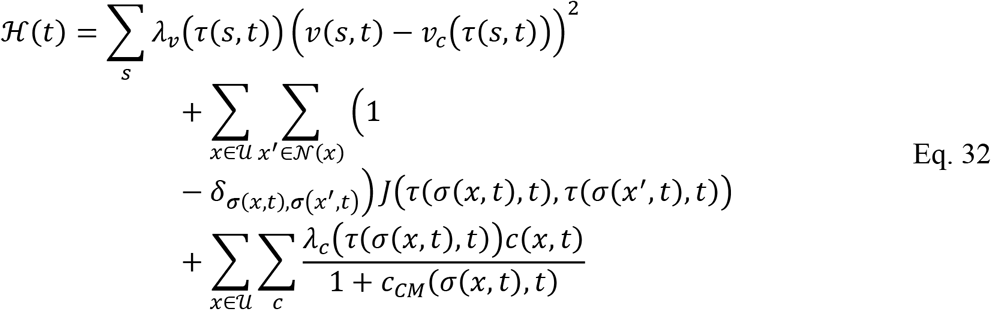

The first summation models a volume constraint for each cell with current volume *v*(*s*, *t*) for cell of identification *s* and type-dependent volume multiplier *λ*_*v*_ and volume constraint *v*_*c*_. The second summation models adhesion, where *δ* is the Kronecker-delta, 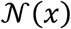 is the neighborhood of a site *x* in the lattice and *j* maps the types occupying the neighboring site pair (*x*, *x*^′^) to a contact coefficient. The third summation models logarithmic chemotaxis due to all simulated fields *c* with type-dependent chemotaxis multiplier *λ*_*c*_ and measurement *c*_*CM*_ of *c* at the center of mass of a cell occupying a lattice site.

In *Results*, we demonstrate cellularization of two ODE models of viral infection in epithelial cells using the spatial configuration employed in (17). The first of the two models represents infection in an epithelial sheet, and the second ODE model adds recruitment of immune cells by inflammatory signaling to the first. As such, epithelial cells are arranged in a uniform, planar configuration and fixed in a regular lattice. For simulations without recruitment of immune cells, the spatial domain is two-dimensional. For simulations with recruitment of immune cells, motile immune cells are placed in a second, two-dimensional layer above the epithelial sheet. Boundary conditions were periodic for boundaries parallel to the plane of the sheet, and Neumann for boundaries perpendicular to the plane of the sheet. The planar dimension and lattice size of all simulations is 400 *μ*m and 200 lattice sizes, respectively (lattice site width of 2 *μ*m/site, Table 1). We applied a uniform volume multiplier of 9 and volume constraint of 25 lattice sites (*λ*_*v*_ and *v*_*c*_ in Eq. 32, respectively) to all cells and placed all epithelial cells in a square shape of 5×5×1 sites. Simulations were performed for two weeks to one month of simulation time, depending on the ODE model and parameter values, with a simulation step of five minutes. All ODE models considered a total number of epithelial cells *N*_*ODE*_ equal to 10 million, from which the global cellularization scaling coefficient *η* was calculated by number of epithelial cells in the spatial domain as *ηN*_*ODE*_ = 1,600 cells, and the local cellularization scaling coefficient *θ* was calculated as *N*_*ODE*_*v*_*c*_*θ* = 1 by imposing the cell volume defined in the spatial model on the epithelial cells described in the ODE models. To test the effects of stochasticity, ten simulation replicas were executed for each spatial model and set of initial conditions and parameters. All simulation measurements were made at intervals of 50 minutes (ten simulation steps). All simulations were performed in CompuCell3D.

**Table 1.** Parameter values used in all simulations.

| Parameter | Value | Reference / Justification |
| --- | --- | --- |
| Simulation step $\Delta t$ | 5 min./step | Selected for approx. 14 days of sim. time in 4k sim. steps |
| Lattice site width | 2 $\mu\text{m}/\text{site}$ | Selected according to cell diameter |
| Volume multiplier $\lambda_v$ | 9 | (17) |
| Volume constraint $v_c$ | 25 sites | Selected according to typical epithelial size and lattice site width |
| Intrinsic random motility $\mathcal{H}^*$ | 10 | (22) |
| Global scaling coefficient $\eta$ | $1.6 \times 10^{-4}$ | Calculated from Eq. 6 according to model specifications |
| Local scaling coefficient $\theta$ | $4 \times 10^{-9} \text{ site}^{-1}$ | Calculated from Eq. 11 according to model specification |

## Results

In this section we demonstrate the cellularization of ODE models of viral infection and immune response according to the formalism defined in *Methods* and deployment of their analogous spatial models in simulation. We present ODE and spatial model simulation results while investigating emergent effects related to spatial and stochastic aspects introduced by the spatial models. Both cellularized ODE models were generated ad-hoc for the purposes of this work according to published models of viral infection and immune response (26,27), the parameters of which were selected only for conveniences of demonstrating their spatial analogues. As such, the simulation results in this work *per se* should not be regarded as relevant to modeling a particular virus or patient scenario. Rather, simulation results should be surveyed within the scope of this work while considering the possible applications of cellularization to existing models of a particular virus or patient scenario.

### Two-dimensional infectivity

The first cellularized ODE model in this work consists of the interactions of a population of cells and an extracellular virus using mass action. The virus infects uninfected cells *U* with an infection rate *β*, which causes uninfected cells to join an infected cell population *I*_1_. Infected cells become virus-releasing cells *I*_2_ at a rate *k*, at which point they release unitless virus *V* at a rate *d*, modeling an eclipse phase. Virus-releasing cells die and join a dead cell population *A* at a rate *d*, modeling virally-induced cell death. The virus decays at a rate *c*,

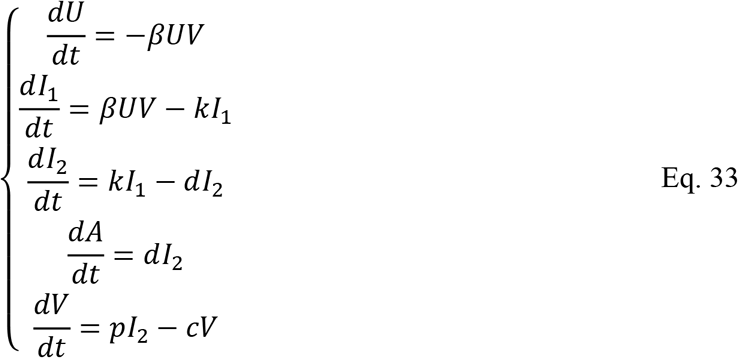

To build a spatial model of local infection using the configuration described in *Implementation Details*, Eq. 33 can be rescaled to a simulated tissue patch and considered as global measurements of the spatial domain. Knowing the total number of epithelial cells in both the ODE and spatial models, the global scaling factor *η* can be calculated from Eq. 6 and applied to generate a form that describes the quantities in Eq. 33 but at the size of the spatial domain,

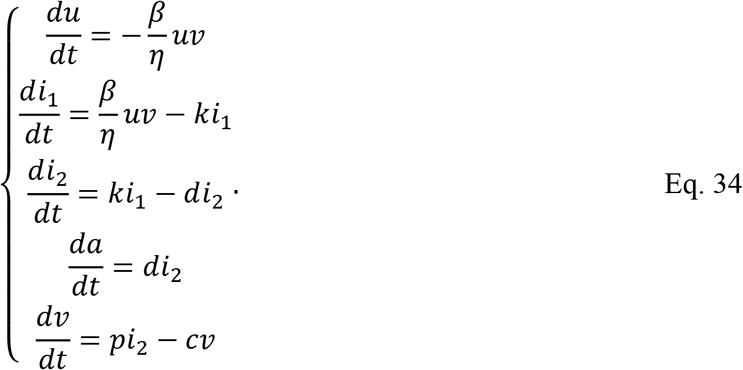

As in *Methods*, here *u* measures the same quantity as *U*, *i*_1_ measures the same quantity as *I*_1_, and so on, but at the scale of the spatial domain. The ODE model describes the processes of virus-mediated infection an uninfected cell 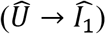, an infected cell becoming a virus-releasing cell 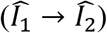, and a virus-releasing cell dying due to the effects of the virus 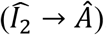. Spatial analogues of these processes can be formulated using Eq. 23 for a diffusive extracellular virus field 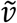 of the form from Eq. 16,

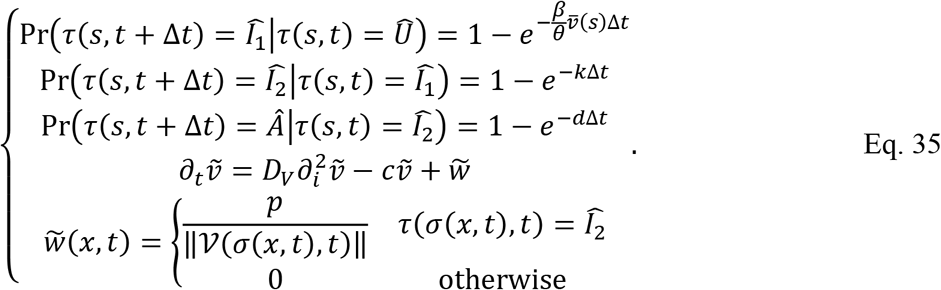

Cellular measurements of extracellular virus 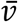 are calculated from 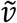 according to Eq. 13. Unless otherwise specified, all simulations were performed with the parameter values in Table 1 and Table 2. Simulations were initialized with fractions of initially infected cells in the range 1% (28) to 10% (29), the distributions of which were random in the spatial model, and with a single initially infected cell, which was placed at the center of the epithelial sheet in the spatial model (17).

**Table 2.** Model parameters used in simulations of infection unless otherwise specified.

| Parameter | Value | Reference / Justification |
| --- | --- | --- |
| Infection rate $\beta$ | $7.18 \times 10^{-13} \text{ s}^{-1}$ | Selected for complete infection by around 5 days |
| Eclipse phase parameter $k$ | $1.85 \times 10^{-5} \text{ s}^{-1}$ | Selected for an eclipse phase of 15 hours |
| Death rate $d$ | $2.78 \times 10^{-6} \text{ s}^{-1}$ | Selected for complete cell death by around 21 days |
| Virus diffusion parameter $D_V$ | $0.1 \mu\text{m}^2 \text{ s}^{-1}$ | Selected to be 10x greater than value from (17) |
| Virus decay rate $c$ | $1.09 \times 10^{-4} \text{ s}^{-1}$ | Selected for a diffusion length of approximately three cells |
| Virus production rate $p$ | $5.79 \times 10^{-4} \text{ cell}^{-1} \text{ s}^{-1}$ | Selected for complete infection by around 5 days |

The distribution of infected and virus-releasing cells demonstrated notable spatial features early in simulation, and especially for fewer initially infected cells (Figure 1). Groups of infected and virus-releasing epithelial cells typically formed near sites of initial infection, which generated prominent gradients in the distribution of extracellular virus. However, some sites of significant infection were also observed early where cells were not initially infected (*e.g.*, Figure 1, 0.01 initial infection fraction, 3 days in simulation time). These sites of infection coalesced into infection throughout the simulated tissue patch, the earliest infected cells of which often, but not always, corresponded to outgrowth of dead cells. Without any anti-viral strategy, nearly all cells died by two weeks of simulation time for all initial infection fractions.

**Figure 1.**
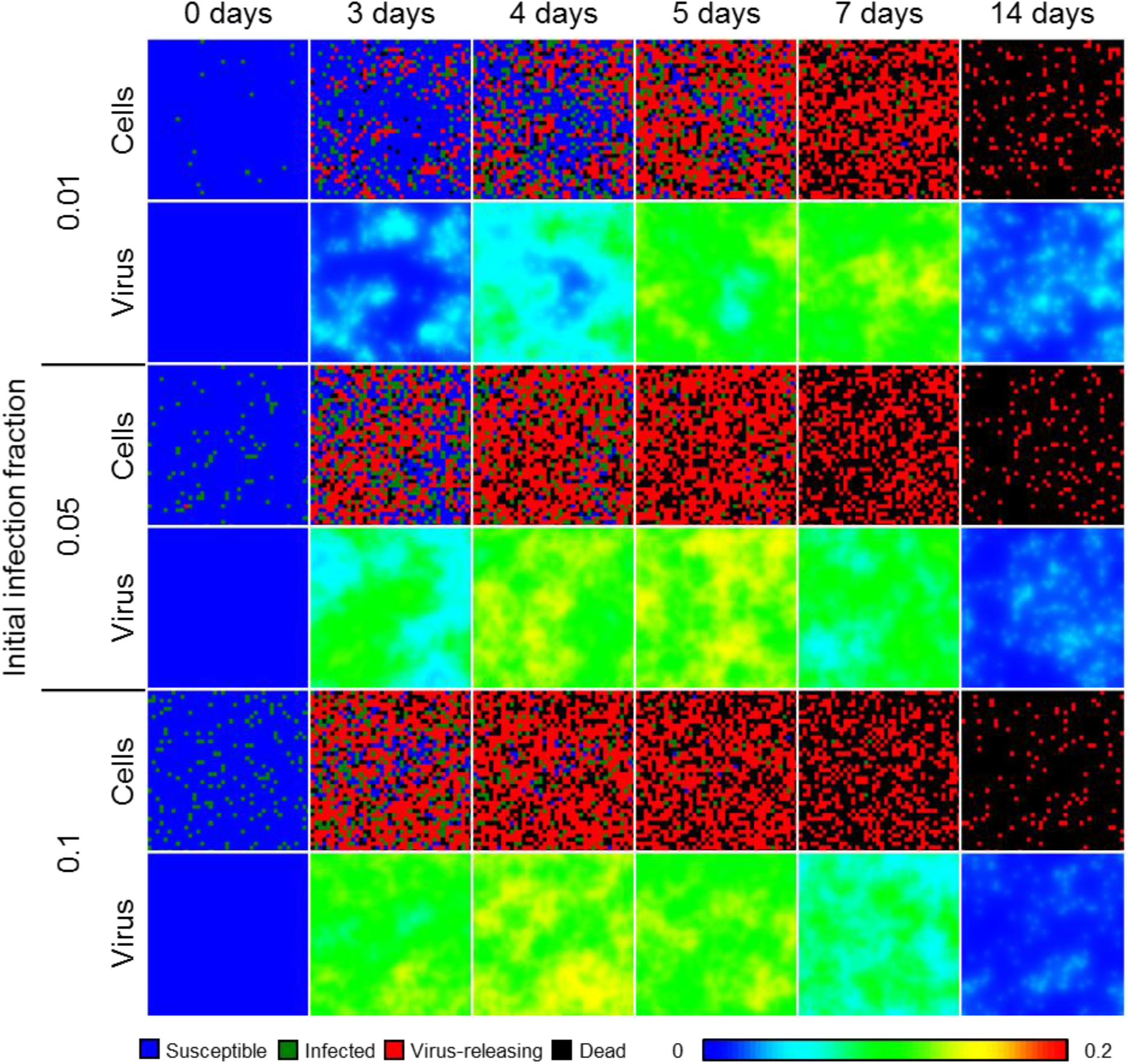
Spatial model results of viral infection in a two-dimensional, epithelial sheet. Results shown for 1% (top), 5% (middle) and 10% (bottom) initially infected cells at 0, 3, 4, 5, 7 and 14 days in simulation time. Epithelial cells shown as blue when susceptible, green when infected, red when virus releasing, and black when dead.

To test the validity of cellularization and the effects of spatial mechanisms introduced to the ODE model, scalar measurements of spatial model results were made of all cell type populations and the extracellular virus and compared to results from the scaled ODE model (Eq. 34). By inspection, the spatial model and employed model parameters generated spatiotemporal dynamics consistently with the original ODE model (Figure 2). Among the ten simulation replicas of the spatial model for each initial infection fraction, only an initial infection fraction of 0.01 generated results with even marginal differences compared those from the scaled ODE model. In the case of an initial infection fraction of 0.01, some simulation replicas produced infection dynamics with a minor delay, where all measures of progression of infection occurred at the same magnitude and with the same dynamical features, but slightly later than predicted by the ODE model.

**Figure 2.**
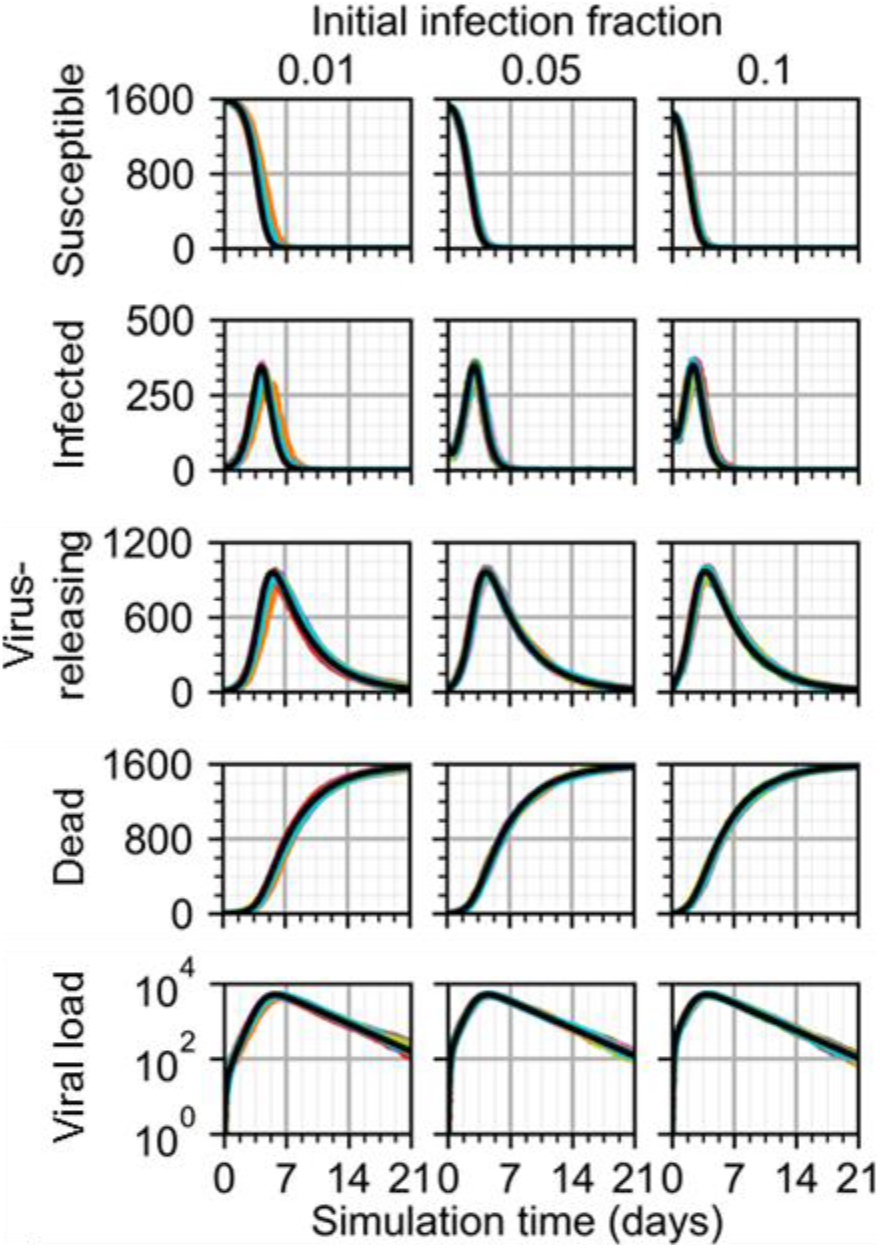
Scalar results from ten replicas of the spatial model of viral infection. Results shown for 1% (left), 5% (center) and 10% (right) initially infected cells. ODE model results shown as black lines.

Spatial effects were even more prominent for simulation replicas with one initially infected cell (Figure 3). Spread of infection exhibited somewhat diffusive, outward spread from the initial site of infection, but with virus-releasing cells far from the initial site of infection even by three days of simulation time (Figure 3A). Such non-diffusive spread of infection was especially noticeable by seven days of simulation, where the distributions of dead cells was comparably distributed as those for an initial infection fraction of 0.1, though the degree of death occurred later in simulation. The observed delay due to lower initial infection was increasingly notable for simulation replicas with one initially infected cell, where peak viral load occurred for some simulation replicas on the order of days later than when predicted by the ODE model (Figure 3B). For one of ten simulation replicas with one initially infected cell, progression of infection did not occur at all.

**Figure 3.**
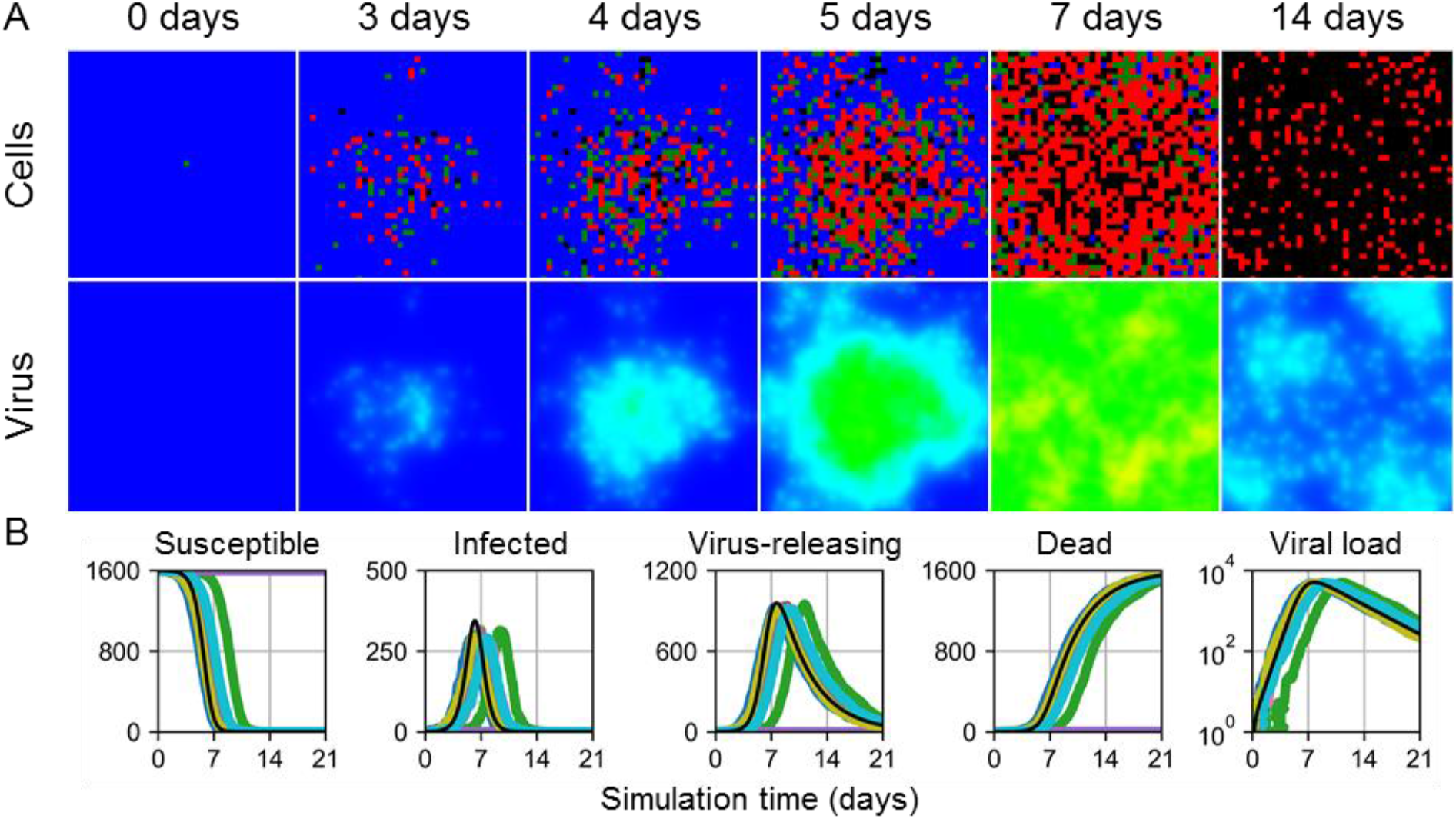
Spatial and ODE model results of viral infection in a two-dimensional, epithelial sheet with one initially infected cell. A. Spatial model results of one simulation replica at 0, 3, 4, 5, 7 and 14 days in simulation time. Epithelial cells and extracellular virus shown as in Figure 1. B. Scalar results from ten replicas of the spatial model. ODE model results shown as black lines.

### Effects of Diffusivity

To test the effects of extracellular diffusion of infection dynamics, the virus diffusion coefficient was swept for all aforementioned initial infection conditions with logarithmic variations over a range of a reduction by a factor of 1,000 from the value in Table 2 (Figure 4). Decreases in virus diffusivity generally led to reductions in the rate of infection of the tissue patch and greater departure from the dynamics described by the ODE model. As in results shown in Figure 3, no infection occurred for some replicas with one initially infected cell. In cases with the minimum considered virus diffusion coefficient (*i.e.*, 0.0001 *μ*m^2^/s), the number of susceptible cells tended towards a non-zero final value. For initial infection fractions of 0.05 and 0.1, the spatial and ODE models agreed for virus diffusion coefficients as low as 0.01 *μ*m^2^/s, whereas the same was true for an initial infection fraction of 0.01 and virus diffusion coefficient as low as 0.05 *μ*m^2^/s.

**Figure 4.**
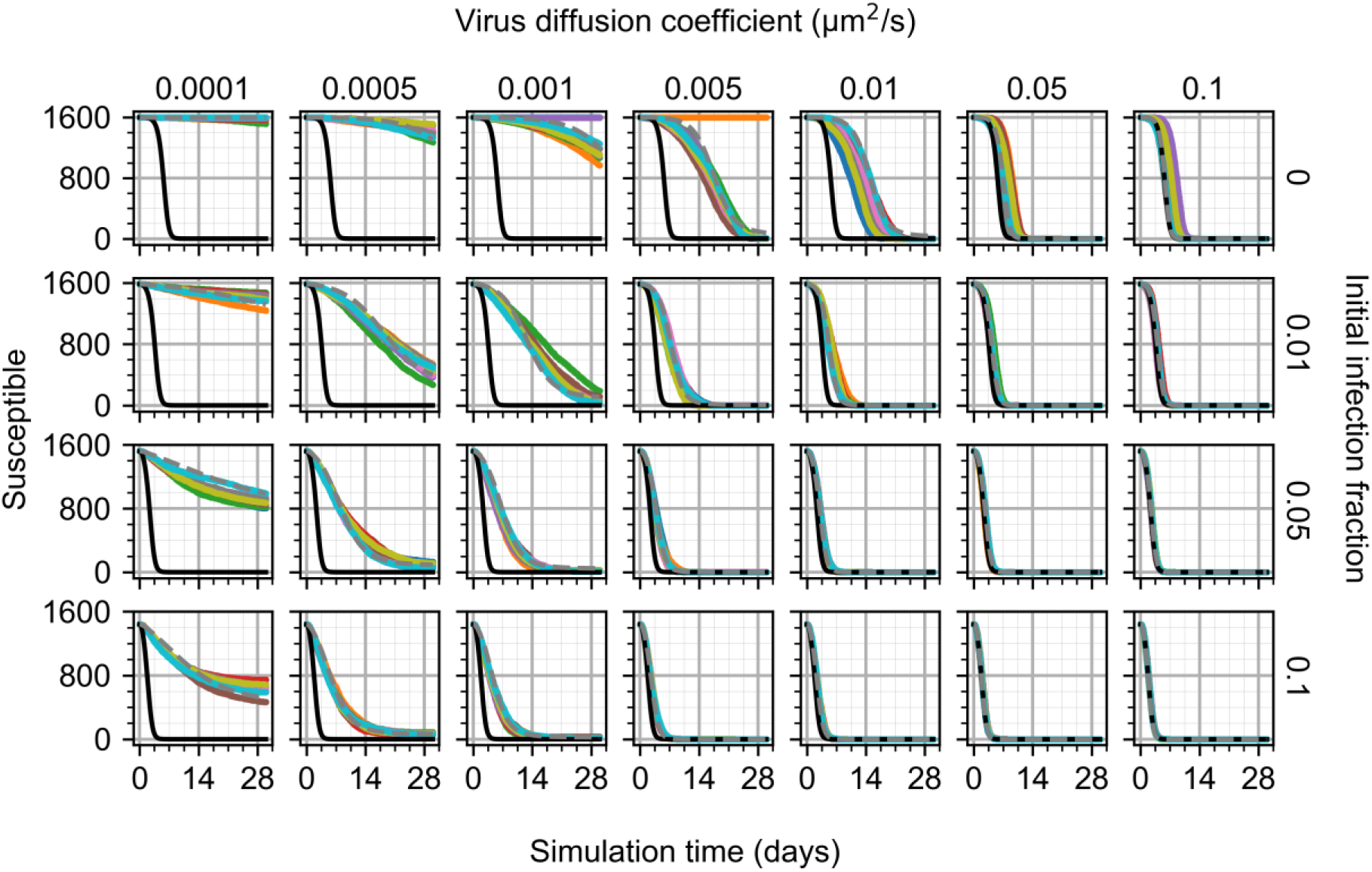
Number of susceptible cells from ten replicas of the spatial model of viral infection while varying initial infection fraction and virus diffusion coefficient. ODE model results shown as black lines. ODE model results with best-fit infectivity shown as gray, dashed lines. Results from the replica used during fitting shown as cyan.

We tested whether decreases in virus diffusion coefficient in the spatial model correspond to decreases in infection rate (*i.e.*, the parameter *β*) by fitting the ODE model to spatial model results while varying the infection rate. Fitting was performed using results from one simulation replica per parameter set and initial conditions when infection occurred. The error during fitting was calculated as the squared difference between the fitted ODE model results and spatial model results for all cell type populations and the viral load at all time points. Reduced infection rates were found that reproduced spatial model results using the ODE model for all cell type populations and viral load (Figure 5) for many virus diffusion coefficients. The ODE model better reproduced spatial model results with reduced infection rates for greater virus diffusion coefficients and initial infection fractions. Some virus diffusion coefficients generated dynamics that were not consistent with the ODE model and reduced infection rates, such as virus diffusion coefficients less than 0.005 *μ*m^2^/s for an initial infection fraction of 0.01, and less than 0.001 *μ*m^2^/s for initial infection fractions of 0.05 and 0.1. Otherwise, the ODE model could be well-fitted to spatial model results with decreasing virus diffusion coefficient by reducing the ODE model infection rate.

**Figure 5.**
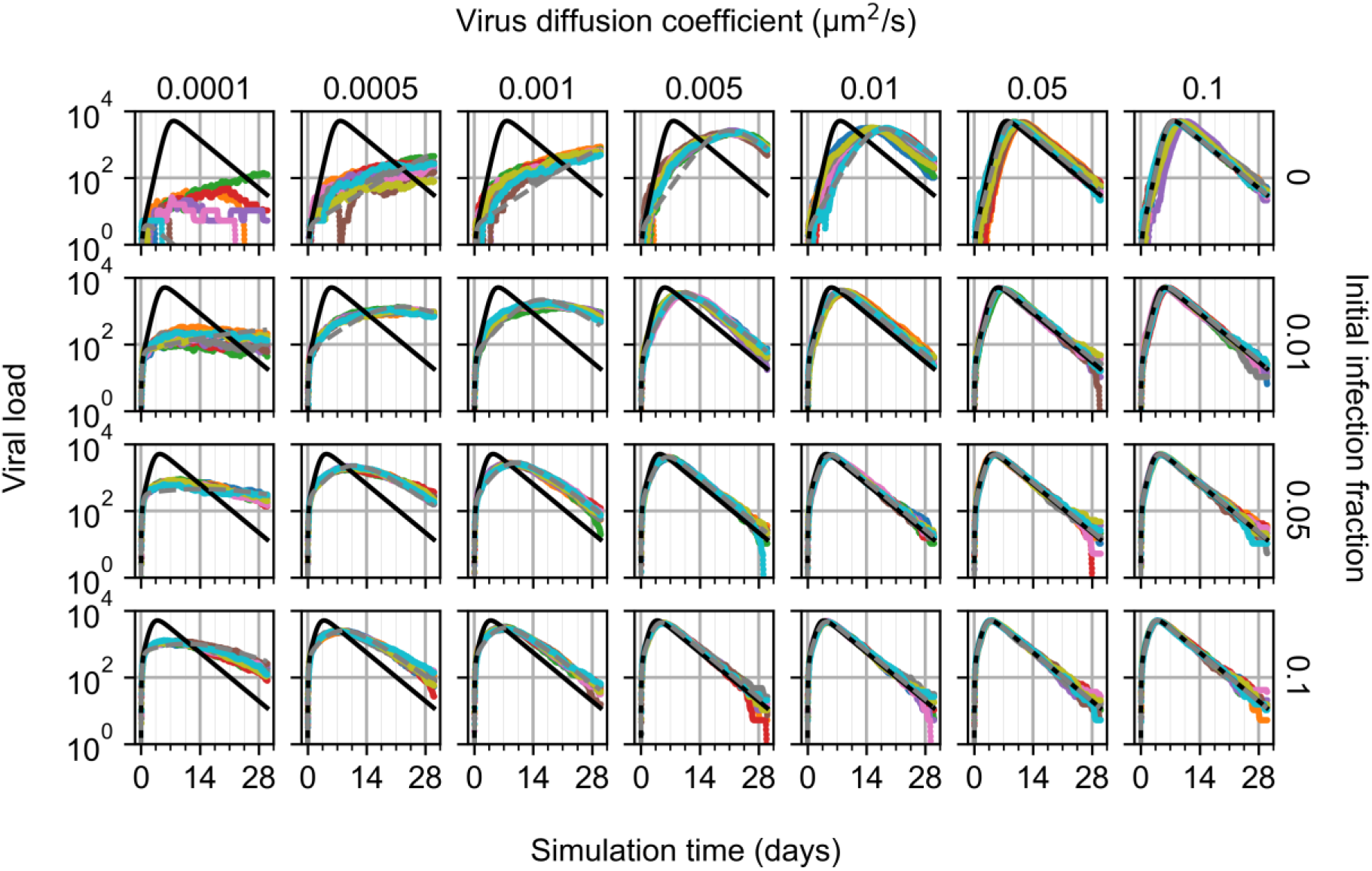
Viral load from ten replicas of the spatial model of viral infection while varying initial infection fraction and virus diffusion coefficient. ODE model results shown as black lines. ODE model results with best-fit infectivity shown as gray, dashed lines. Results from the replica used during fitting shown as cyan.

### Immune response modeling

The second cellularized ODE model in this work adds a compartmental model of immune cell proliferation and recruitment to the ODE model of viral infection in *Two-dimensional infectivity*. Infected and virus-releasing cells release a local cytokine *C* at a rate *p*_*C*_, which decays at a rate *c*_*C*_. The local cytokine transports to cytokine in a lymph node compartment *C*_*L*_ at a rate *k*_*C*_, which decays at a rate *c*_*cl*_. The lymph node cytokine induces proliferation of an immune cell population in the lymph node compartment *E*_*L*_ at a rate *p*_*el*_, which also undergoes saturated proliferation and transports to a local immune cell population *E* at a rate *k*_*e*_. Each local immune cell kills virus-releasing cells at a rate *d*_*ei*2_ (26), and the local immune cell population decays at a rate *d*_*E*_,

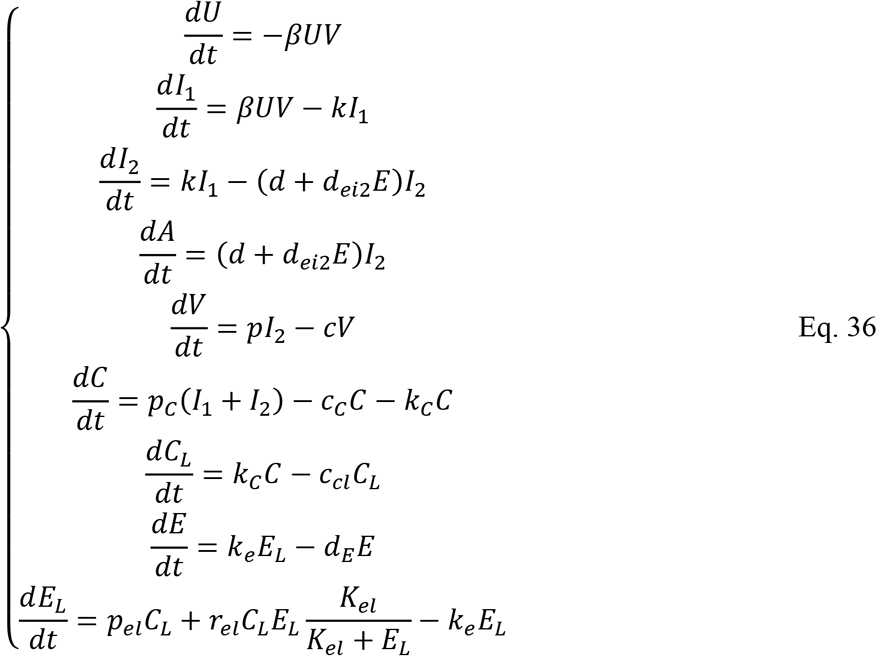

Continuing the construction of a spatial model of a local patch of infection as described in *Two-dimensional infectivity,*Eq. 36 can be scaled in the same manner as performed on Eq. 33 using the total number of epithelial cells in the spatial and ODE models,

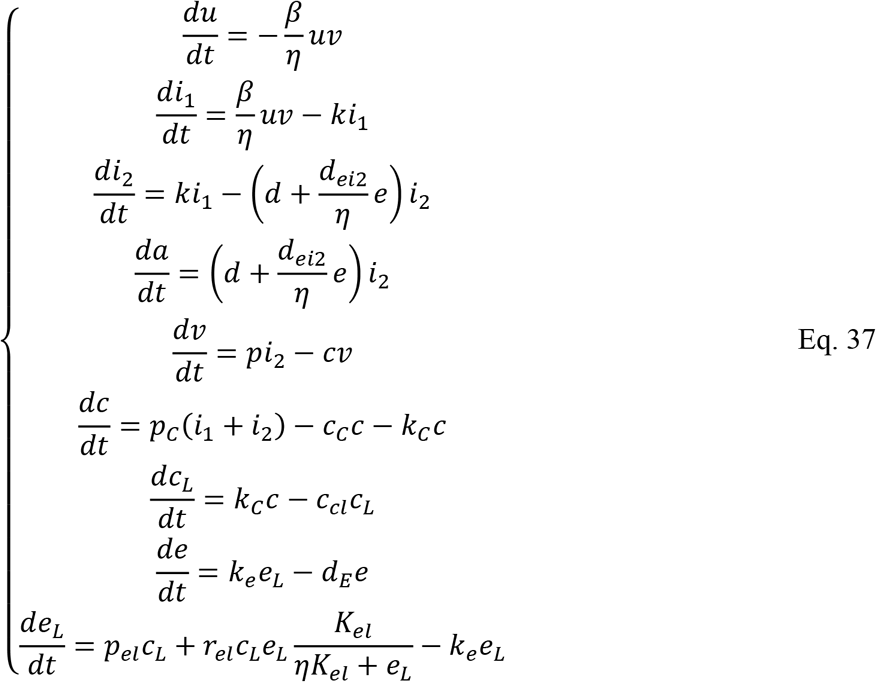

Note that rescaling the cytokine and immune cell population in the lymph node compartment is optional, and could instead remain at the scale of the original ODE model (*e.g.*, *e*_*L*_ = *ηE*_*L*_ throughout). In the spatial model, local cytokine and each local immune cell are spatially modeled, whereas the lymph node cytokine and immune cell population remain as ODEs. Contact-mediated killing is cellularized for each virus-releasing cell using Eq. 28 for 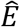 − type (*i.e.*, local immune cell type) cell total contact area *A*_*E*_, total number of epithelial cells *N*_*e*_, and total contact area between a cell *s* and 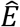 cells *A*_*σ*,*s*_. As such, Eq. 36 can then be written according to the aforementioned multiscale structure,

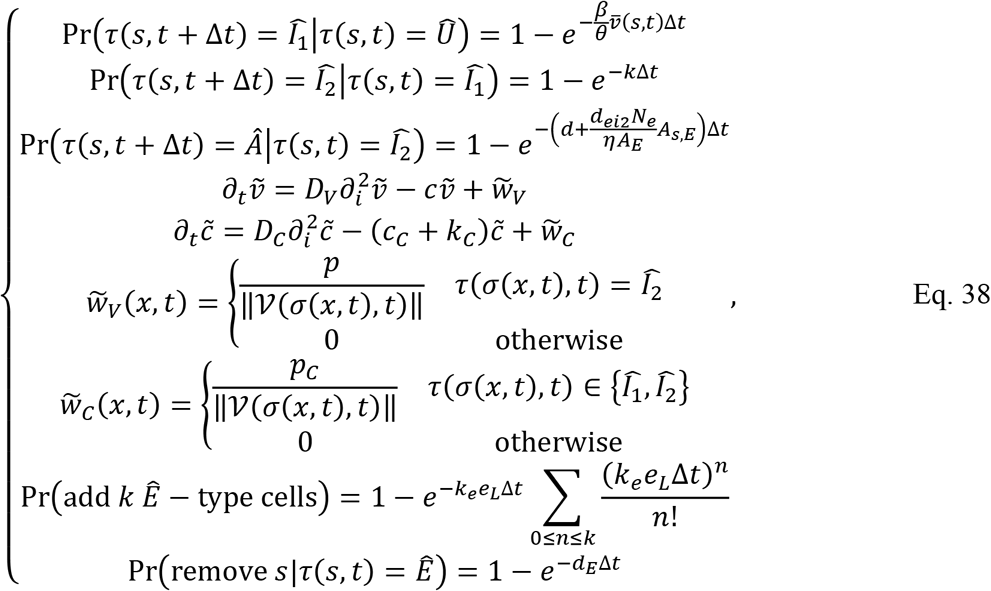

where the rate equations for *e*_*L*_ and *c*_*L*_ are the same as in Eq. 37. All other equations are described in *Two-dimensional infectivity*. The spatial domain boundary above the epithelial sheet was assigned as the inflow boundary for local immune cells. Outgoing cells were immediately removed from the spatial domain and replaced with medium. Unless otherwise specified, all simulations were performed with the parameter values in Table 1, Table 2 and Table 3.

**Table 3.** Model parameters used in simulations of infection and immune response unless otherwise specified.

| Parameter | Value | Reference / Justification |
| --- | --- | --- |
| Immune killing rate $d_{ei2}$ | $1.16 \times 10^{-11} \text{ cell}^{-1} \text{ s}^{-1}$ | Selected for comparable killing rate by viral death |
| Cytokine production rate $p_C$ | $1.16 \times 10^{-4} \text{ cell}^{-1} \text{ s}^{-1}$ | Selected for maximum total cytokine on the order of 1k in typical simulations |
| Cytokine decay rate $c_C$ | $2.31 \times 10^{-5} \text{ s}^{-1}$ | Selected for a diffusion length of approximately eight cells |
| Cytokine transport rate $k_C$ | $5.79 \times 10^{-6} \text{ site s}^{-1}$ | Selected for appreciable delay in inflammatory signaling |
| Lymph node cytokine decay rate $c_{cl}$ | $5.79 \times 10^{-6} \text{ s}^{-1}$ | Selected for cytokine clearance by two weeks |
| Immune cell transport rate $k_e$ | $1.16 \times 10^{-5} \text{ s}^{-1}$ | Selected for appreciable delay in immune cell recruitment |
| Immune cell decay rate $d_E$ | $1.16 \times 10^{-5} \text{ s}^{-1}$ | Selected for immune cell clearance by two weeks |
| Cytokine-induced immune cell production rate $p_{el}$ | $1.16 \times 10^{-9} \text{ cell s}^{-1}$ | Selected for marginal induction of immune cell production |
| Cytokine-activated immune cell production rate $r_{el}$ | $5.79 \times 10^{-8} \text{ s}^{-1}$ | Selected for significant rate-limited production of immune cell production |
| Immune cell production midpoint $K_{el}$ | 500 cells | Selected for maximum total immune cells on the order of 3k in typical simulations |
| Number of local epithelial cells $N_e$ | 1,600 cells | Calculated from spatial model specification |
| Total contact surface area $A_E$ | 25 site surfaces | Assumed based on epithelial sheet geometry |
| Cytokine diffusion parameter $D_C$ | $0.16 \mu\text{m}^2 \text{ s}^{-1}$ | (17) |
| Seeding fraction | 0.1% | Selected for mostly random immune cell seeding |
| Chemotaxis multiplier $\lambda_c$ | 10,000 | Typical CPM value for reasonable chemotaxis |
| Immune cell-medium contact coefficient | 20 | Selected for preferential attachment of immune cells not with each other |
| All other contact coefficients | 25 | Selected for preferential attachment of immune cells not with each other |

The immune response prevented complete spread of infection throughout the spatial domain in all ten simulation replicas of 0.01, 0.05 and 0.1 initial infection fraction, and decreasingly so with increasing initial infection fraction (Figure 6). After two days of simulation time, infected and virus-releasing cells were more prominently distributed throughout the spatial domain compared to local immune cells. However, a strong immune response recruited sufficient immune cells to nearly cover the entire epithelial sheet by days 7, 6 and 5 for initial infection fractions 0.01, 0.05 and 0.1, respectively, resulting in significant killing of virus-releasing cells and prevention of further infection. By two weeks of simulation time for all initial conditions and simulation replicas, virus and cytokine levels were near zero, most immune cells had left the spatial domain and the epithelial sheet consisted of susceptible cells with significant distributions of dead cells. Initial infection fractions of 0.01, 0.05 and 0.1 resulted in final fractions of dead cells of around 0.5, 0.625 and 0.75, respectively.

**Figure 6.**
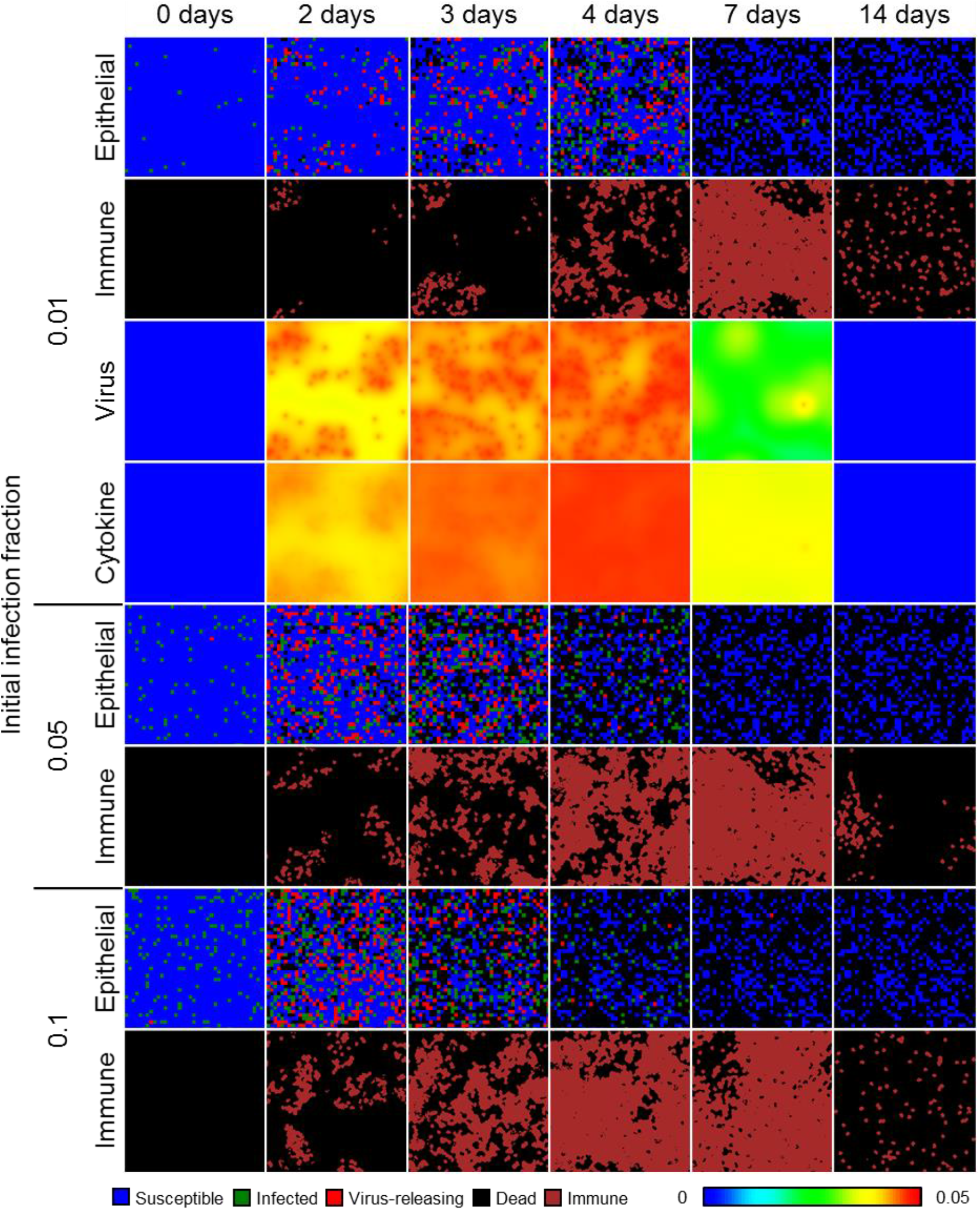
Spatial model results of viral infection and immune response in a quasi-two-dimensional, epithelial sheet. Results shown for 1% (top), 5% (middle) and 10% (bottom) initially infected cells at 0, 2, 3, 4, 7 and 14 days in simulation time. Epithelial cells shown as in Figure 1. Immune cells shown as dark red.

As with the model of viral infection described in *Two-dimensional infectivity*, the spatial model of viral infection and immune response generated spatiotemporal dynamics consistently with the original ODE model using the employed model parameters (Figure 7). Simulation replicas of the spatial model produced nearly the same number of susceptible cells at the end of simulation as the ODE model for 0.01 initial infection fraction, while replicas of the spatial model produced slightly fewer final susceptible cells and more final dead cells than the ODE model for initial infection fractions of 0.05 and 0.1. For an initial infection fraction of 0.01, peak viral load, local cytokine and lymph node cytokine in replicas using the spatial model occurred slightly later than for the ODE model, while these metrics were approximately the same between the spatial and ODE models for initial infection fractions of 0.05 and 0.1. The number of infected, virus-releasing and lymph immune cells significantly varied for replicas of the spatial model and an initial fraction of 0.01, while for initial infection fractions of 0.05 and 0.1 the maximum number of infected, virus-releasing and lymph immune cells were slightly greater than those of the ODE model. However, spatial model replicas produced slightly fewer local immune cells than the ODE model before peak viral load and then tended towards the same values as the ODE model thereafter.

**Figure 7.**
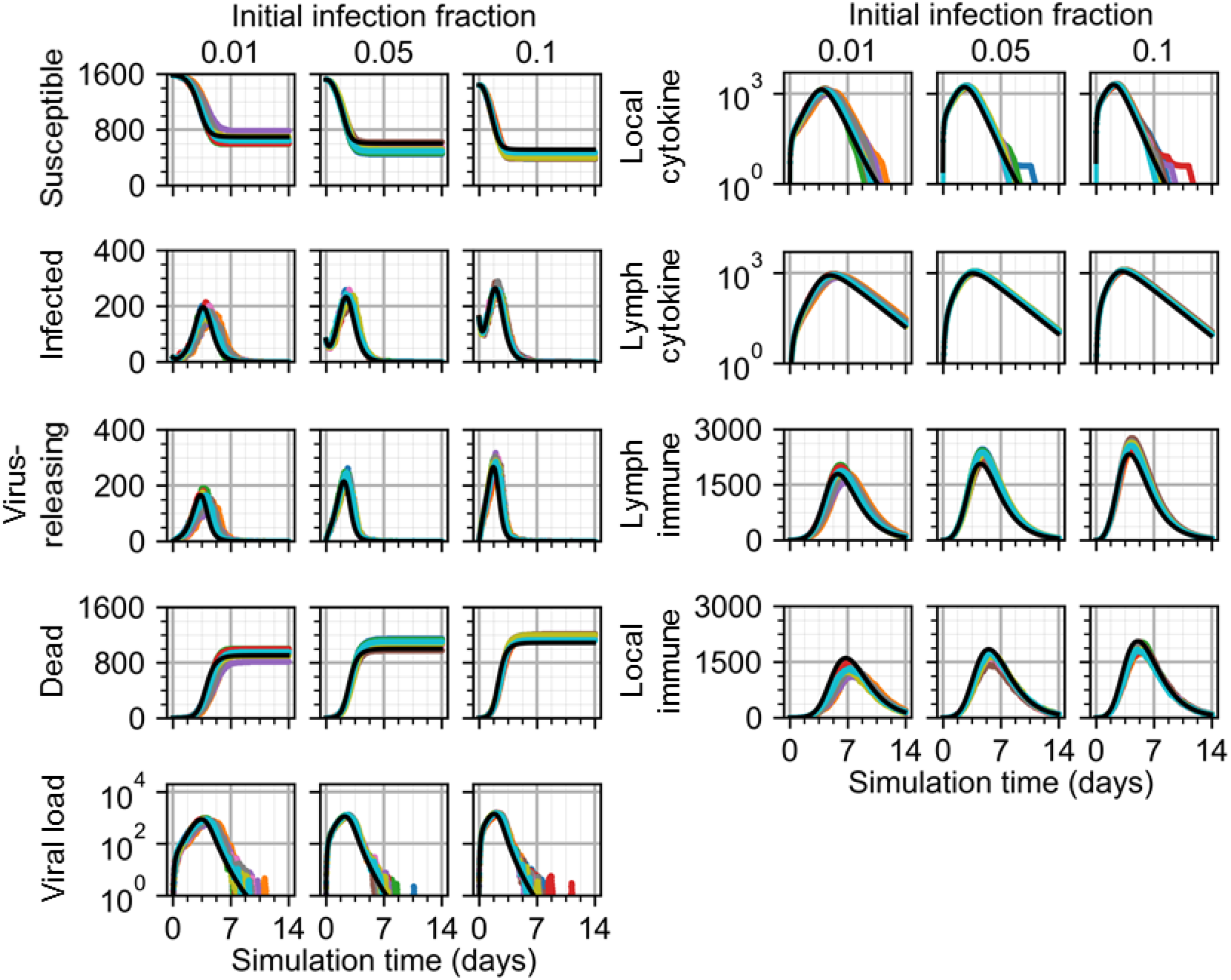
Scalar results from ten replicas of the spatial model of viral infection and immune response. Results shown for 1% (left), 5% (center) and 10% (right) initially infected cells. ODE model results shown as black lines.

In some simulation replicas of the spatial model, viral load suddenly increased to appreciable values even days after viral load decreased below a value of 1, and then decreased again to negligible values. Inspection of simulation data showed that latent extracellular virus infected susceptible cells after the initial progression of infection in some replicas of the spatial model. We observed this event, which cannot occur using the ODE model, in previous work modeling viral infection and called it “recursion” (17).

We tested the effects of spatial mechanisms introduced during development of the spatial analogue to the ODE model of infection and immune response. Specifically, we performed a parameter sweep of the sampling fraction by which immune cells are seeded and the chemotaxis parameter that describes how sensitively they chemotax along gradients of local cytokine for an initial infection fraction of 0.1. We varied the sampling fraction from 2.5×10^−5^ (*i.e.*, equivalent to seeding in random locations) to 1.0 (*i.e.*, at the maximum value of local cytokine in all available sites), and the chemotaxis parameter from 10^4^ (*i.e.*, virtually no chemotaxis) to 10^6^ (*i.e.*, just below permitting annihilation of cells by excessive chemotactic forces) using a logarithmic scale. We measured the normalized root mean squared error (NRMSE) of results from each of ten simulation replicas as forecasting results from the ODE model. NRMSE calculations were made for results in intervals of ten simulation steps (*i.e.*, 50 minutes) for each parameter set, the summation of which we used to quantify how well the spatial model represented the constituent components of the ODE model using each parameter set.

As determined by the final number of susceptible cells in simulation replicas, we found that the most effective immune response strategy was one of random seeding and strong chemotaxis, and the least effective strategy was one of cytokine-mediated seeding and marginal chemotaxis (Figure 8). Seeding cells at the currently available maximum (*i.e.*, a seeding fraction of 1) produced the least effective immune response to prevent infection for a given chemotaxis parameter, as was marginal chemotaxis for a given sampling fraction. Likewise, seeding immune cells mostly at random produced a better strategy for mitigating spread of infection as did stronger chemotactic sensitivity for a given sampling fraction. Immune response was most effective for sampling fractions up to 0.1%, and was more effective with increasing sampling fraction and decreasing chemotaxis parameter.

**Figure 8.**
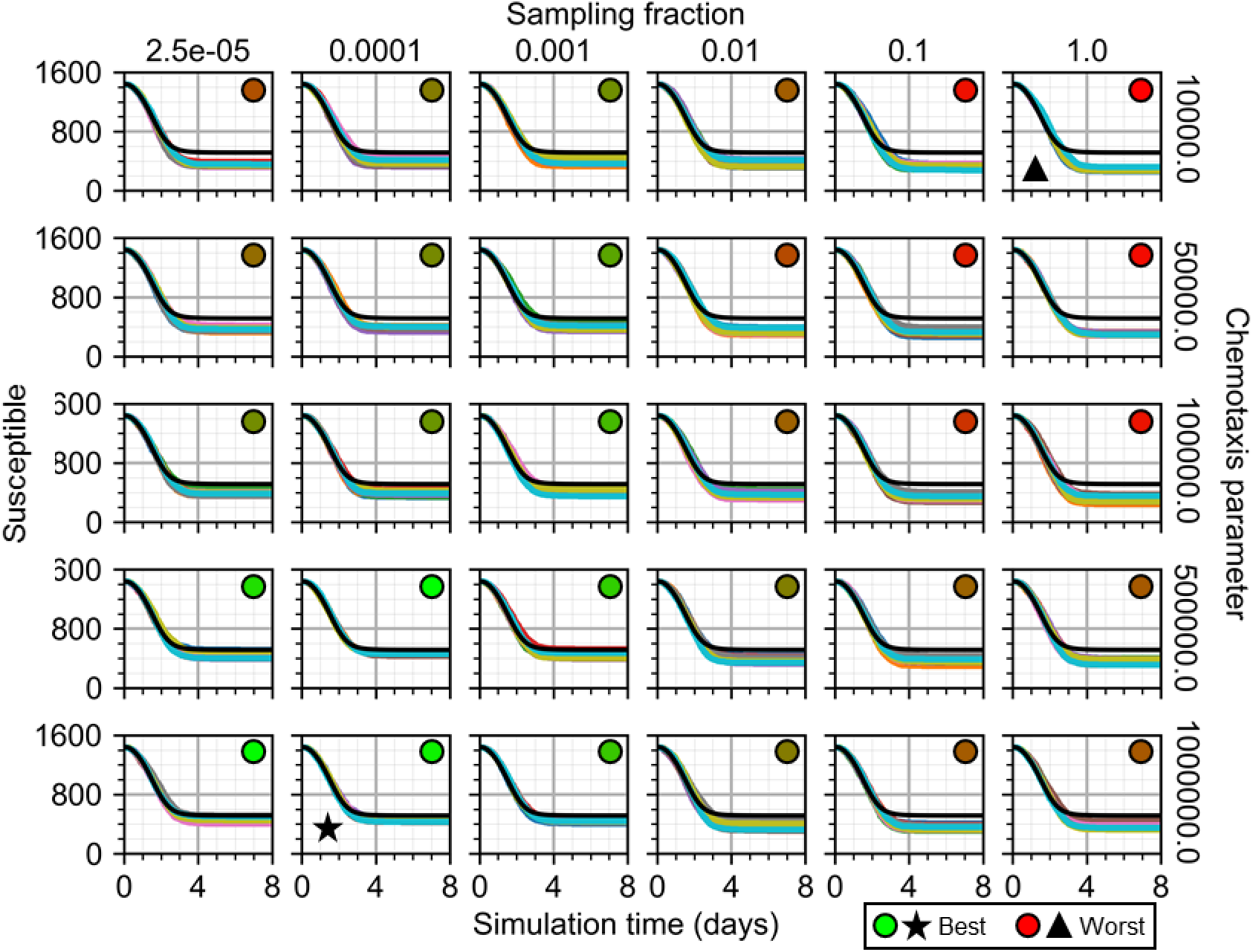
Susceptible cells from ten replicas of the spatial model of viral infection while varying immune cell sampling fraction and chemotaxis parameter. ODE model results shown as black lines. Star marks best. Triangle marks worst. Circles with green to red shading show best to worst total error, respectively.

A significant source of differences between the spatial and ODE models was differences in the timing of peak viral load during progression of infection (Figure 9). For greater sampling fractions and lesser chemotaxis parameters, peak viral load was greater and occurred later in replicas using the spatial model. These differences produced viral load curves for simulation replicas of the spatial model with the same dynamical features as the ODE model (maximum value around two to three days, followed by logarithmic decay to zero), and even (nearly) identical results before peak viral load, but with continued progression of infection up to a day longer (*e.g.*, sampling fraction of 1.0 and chemotaxis parameter of 10^4^) before decline and elimination of infection. For mostly random seeding and strong chemotaxis (*e.g.*, sampling fraction of 2.5×10^−5^ and chemotaxis parameter of 10^6^), the only notable differences between spatial and ODE model results occurred long after peak viral load and for all parameter sets, where single infection events produced strong fluctuations in otherwise negligible viral load values. The overall best-fit parameter set was a sampling fraction of 0.01% and chemotaxis parameter of 10^6^.

**Figure 9.**
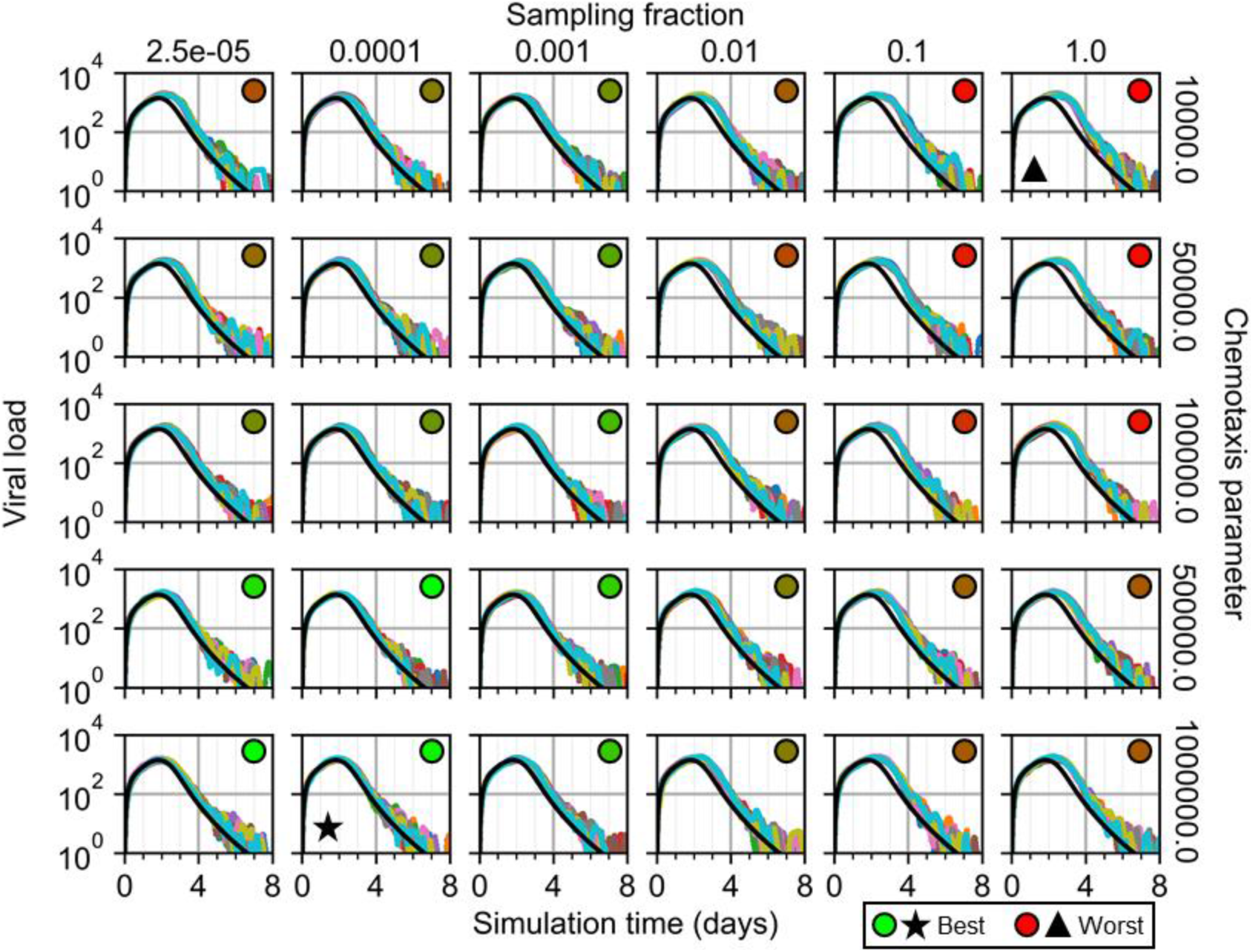
Viral load from ten replicas of the spatial model of viral infection while varying immune cell sampling fraction and chemotaxis parameter. ODE model results shown as black lines. Star marks best. Triangle marks worst. Circles with green to red shading show best to worst total error, respectively.

Lastly, we tested the best-fit parameters for aforementioned parameter sweep for the scenario of one initially infected cell. Previously observed stochasticity of system dynamics and outcomes for one initially infected cells in *Two-dimensional infectivity* were also observed when using the best-fit parameters (Figure 10). In the case of also simulating an immune response, the final distribution of local immune cells varied significantly among ten simulation replicas. In simulation replicas that experienced more infection, few immune cells were found near the site of initial infection, whereas in replicas that experienced less infection, more immune cells were found nearer to the site of initial infection.

**Figure 10.**
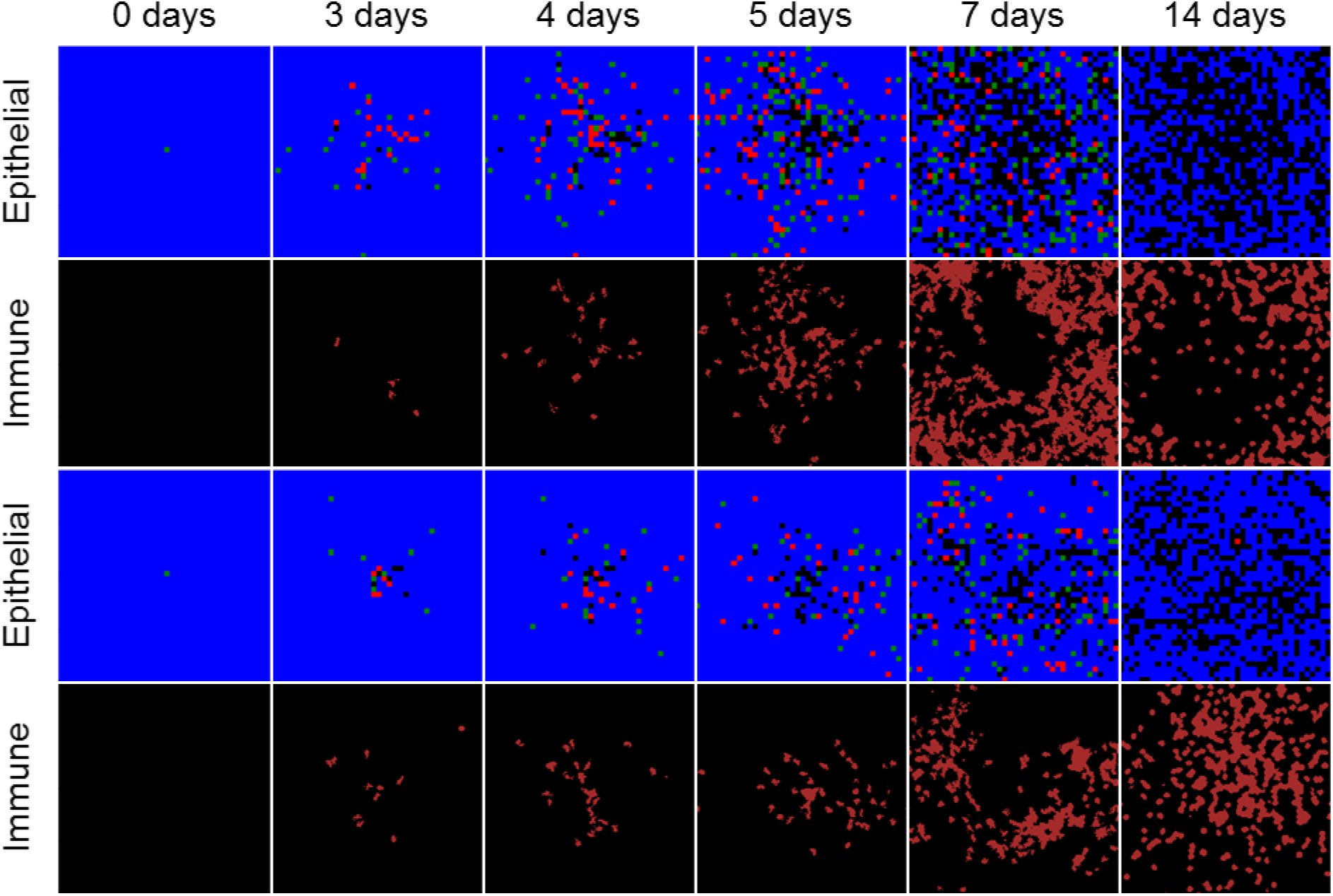
Spatial model results from two simulation replicas of viral infection and immune response in a quasi-two-dimensional, epithelial sheet with best-fit immune response parameters and one initially infected cell. Results shown at 0, 3, 4, 5, 7 and 14 days in simulation time. Epithelial and immune cells shown as in Figure 6.

Two simulation replicas experienced no significant infection, and those replicas that did experienced infection at significantly varying degrees of severity (Figure 11). Among replicas with more than one infected cell during simulation time, the final number of susceptible cells was in the range of 772 to 1,045 cells, with mean and standard deviation of 880 and 98.7 cells, respectively. Compared to a final number of susceptible cells of 722.7 according to the ODE model, the spatial model produced results that disagree with the ODE model in the range of about 7% to 45% less infection for replicas that experienced infection. Measurements of viral load and local and lymph cytokine were comparable to the ODE model in magnitude, though the replicas that experienced less severe infection also experienced later peak viral loads and cytokine, with corresponding effects on immune cell recruitment.

**Figure 11.**
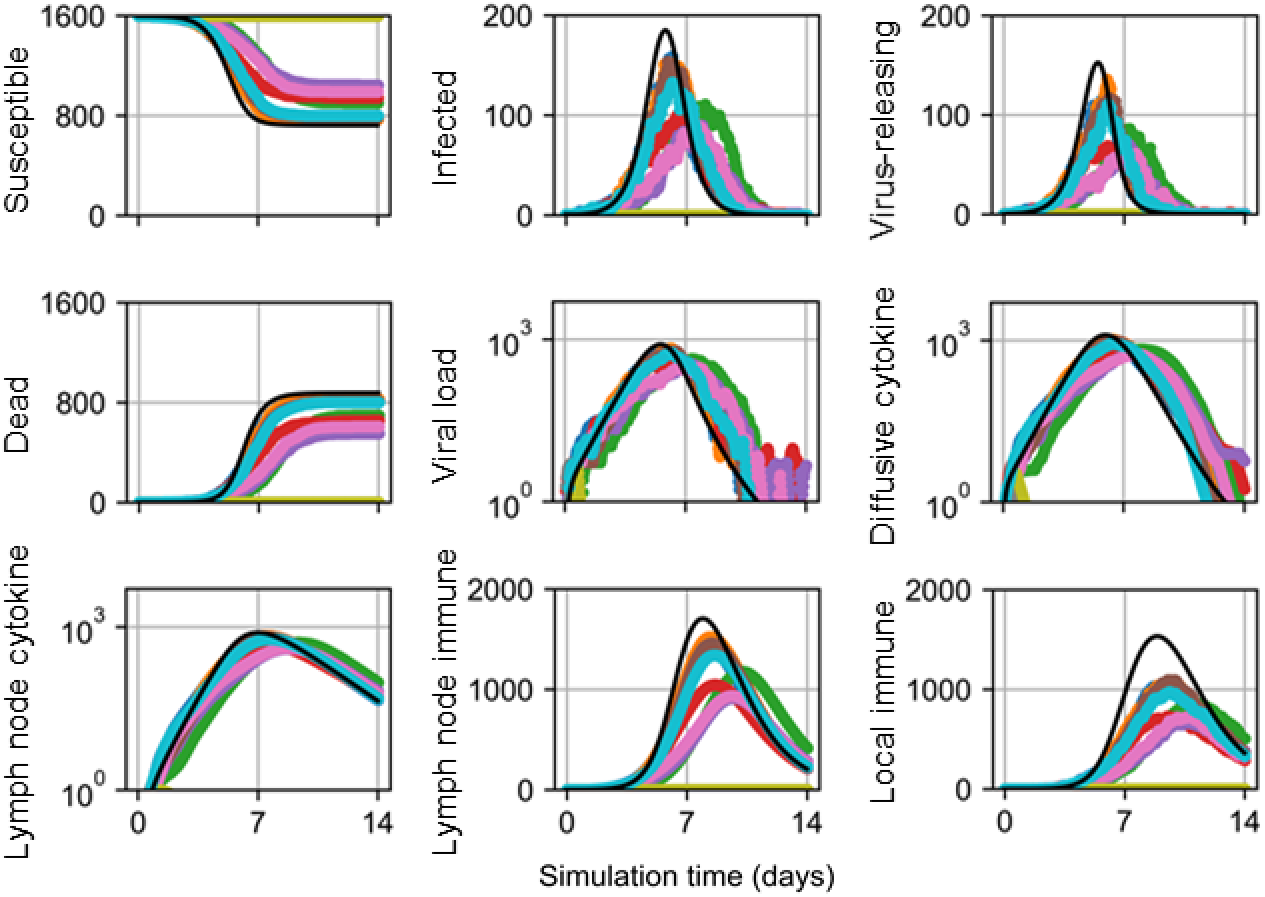
Scalar results from ten replicas of the spatial model of viral infection and immune response with best-fit immune response parameters and one initially infected cell. ODE model results shown as black lines.

## Discussion

We demonstrated that spatial mechanisms like mass transport are implicitly represented in non-spatial model parameters like virus infection rate. This aspect of a non-spatial model is particularly important when considering how to interpret *in vitro* results and translate information gained from *in vitro* scenarios to predictions of *in vivo* outcomes. For example, diffusive transport of extracellular virions during viral infection depends on the medium through which virions migrate, which varies *in vitro* by culture medium and *in vivo* by location. Results in *Effects of Diffusivity* can be regarded in this context as characterizing the effects of the environment on progression of infection while holding all other mechanisms constant, of which we showed to significantly affect both the rate and severity. Such observations then elucidate a means by which we can interrogate when coarse-grained approaches fail to reliably predict physical data by examining the underlying spatiotemporal mechanisms. As demonstrated in this work, our method of cellularization enables such activities related to both synthesis and validation of non-spatial models.

We also demonstrated how generating a spatial analogue of a non-spatial ODE model can further elucidate the features of the underlying mechanisms that a non-spatial model only implicitly represents. The parameter sweep of spatial model parameters in *Immune response modeling* showed the possible strategy of the immune response to eliminate viral infection. Specifically, immune cells more effectively induce apoptosis in infected cells via contact-mediated interactions when they arrive at a mostly random location near a local site of infection and then strongly chemotax along soluble immune signals. Probably, arriving where immune signals are the strongest results in immune cells that cannot effectively prevent the advancing front of an infection, due to the nature of diffusive transport. Rather, it may be more effective to arrive near, but just outside, a lesion. This suggests that resident cells of a site of infection may further activate chemotactic sensitivity in the immune cells described by an ODE model like ours that models viral infection and immune response (e.g., CD8+ T cells). Such observations well describe recruitment and search strategies within the context of current understanding of effector T cells (30). Namely, that effector T cell search strategies for infected cells balance a trade-off of exploration and exploitation (*i.e.*, of random and directional migration), and that T cells presumably encounter antigen-presenting cells much more often in inflamed tissue than in lymph nodes, which changes their search strategy from exploration-dominant to exploitation-dominant.

Some simulation replicas of viral infection generated no infection of tissue even without an immune response for various model parameters and initial conditions. This observation is particularly important with regards to both the biological features represented in the non-spatial and spatial models, as well as the meaning of spatial model, of viral infection. It is certainly a reasonable and relevant question to ask whether a single infected cell can cause infection in tissue, and to what extent. For such a question, the ODE model always predicts infection, due to its continuous treatment of discrete objects and processes (*e.g.*, a half of a virion can generate a quarter of an infected cell in the ODE model), while the spatial analogue allows for premature death and ineffective viral exposure. In this regard, the spatial analogue better describes the total story of infection. This is not to say that either is a sufficiently insightful or predictive model of viral infection *per se*. Rather, we argue that the spatial analogue provides a better description of the biophysical mechanisms responsible for viral infection according to the ODE model. The feasibility of employing such a spatial analogue on useful modeling applications could be questioned due to the greater computational cost of multicellular simulations, though such a limitation is technological. To this end, our cellularization method improves the feasibility of using spatial, multicellular models by readily permitting the employment of model parameters fitted to *in vivo* and/or *in vitro* data using analogous non-spatial models, which are computationally inexpensive.

More importantly, deciding on the context of the spatial domain of a spatial analogue with respect to the entire biological system is a particularly challenging modeling problem. For example, the spatial analogues developed in this work employed periodic boundary conditions on the boundaries orthogonal to the epithelial sheet. This arrangement implies that the spatial analogue describes one instance of a periodic system, the collection of which the ODE model describes. This paradigm breaks down when considering highly localized cases like one initially infected cell, as demonstrated by multiple simulation replicas of the spatial analogue producing significantly different outcomes (*i.e.*, widespread infection or no infection). In such a case, we could observe no infection in one replica and then argue that infection occurred in some other replica, at the expense of revising the premise of the model altogether (*i.e.*, the system described by the ODE model is not periodic). As inconvenient as such a scenario may be, it demonstrates that cellularization better equips the modeler to interrogate the underlying mechanisms responsible for the emergent dynamics of a biological system. In the case of viral infection, cellularization enables the utilization of both well-fitted ODE models of *in vivo* data and *in vitro* evidence of viral transport and discrete viral internalization by inquiring about the cellular, multicellular and spatiotemporal mechanisms that can consistently explain observations at multiple scales.

### Future Work

In the immune response model, the spatial model produced greater lymph immune cells and fewer local immune cells than the ODE model. These differences may be due in part to that the seeding algorithm always considers increments in the number of cells under consideration for seeding beginning at zero. The significance of this apparent bias is currently unclear, as is any exact mitigation strategy, though it may be possible to refine the seeding algorithm by adding stochasticity to the considered number of seeded cells per algorithmic iteration.

There are many mathematical terms employed by non-spatial models of biological systems to which the general forms described in this work (*e.g.*, Eq. 15, Eq. 20, Eq. 24) cannot be readily applied to generate linear stochastic transition rules during cellularization (*e.g.*, Michaelis-Menten). Deriving spatial analogues of ODE model terms not only provides broader utility by allowing the development of an analogous spatial model, but also a stronger understanding of the mechanism(s) represented by the ODE model term. Cellularization of Eq. 33 provided a very simple example of such a case, where casting the infectivity *β* in cellular terms demonstrates it as an extensive property, which limits its interpretation without additional knowledge of the described system (as opposed to an intensive infectivity, which explicitly describes the interactions of a particular cell population and virus). Likewise, proving the nonexistence of a spatial analogue calls into question what exactly an ODE model term means. In such a case for an ODE model term that describes a cellular interaction, questions arise as to what a cellular model means if it cannot be written on a cellular basis. This is not to propose that the validity of an ODE model depends on its mathematical compatibility with cellularization. Indeed, many ODE models have been well-fit to experimental data, some terms of which a spatial analogue is not obvious to us (26,31). Rather, we propose that an ODE model term that is incompatible with cellularization represents multiple mechanisms, each of which can be described by an ODE model term that supports cellularization. The justification for this proposition is obvious from a biological perspective, in that any subsystem of an organism defined at a scale higher than the cellular scale can be resolved to a cellular basis, which is the basis of spatial models according to our cellularization method. Beyond providing a clear path to generating spatial analogues, the validity of this proposition is relevant to many fields related to mathematical biology. A clear separation of mechanisms provides further insights into the dynamics of a biological system relevant to both biomedical sciences (*e.g.*, when developing drug therapies) and basic biology (*e.g.*, when quantitating biological complexity), as well as a more descriptive and robust mathematical model. As such, we propose that the mathematical compatibility of an ODE model with cellularization is an indicator of the robustness of the ODE model.

We can also turn this paradigm—of regarding ODE models within the context of generating a spatial analogue—towards spatial models of multicellular systems while considering their non-spatial analogue. Indeed, previous works have performed related investigations of how spatial and cellular mechanisms affect emergent dynamics in biological systems of which non-spatial models have been developed. One such example described emergent growth dynamics in diffusion-limited systems on a cellular basis as affected by aggregate shape (16), which could be further investigated with regards to existing non-spatial models of organoid growth *in vitro* (32,33) using cellularization. Furthermore, operations defined in our cellularization method on spatial models may generate novel non-spatial model terms that are not obvious from a homogenized approach. We are currently investigating such an approach for describing cell-specific intracellular viral replication dynamics and cell- and position-specific cytotoxic killing, in the fashion of our recent work modeling viral infection and immune response (17). This approach may also be useful for introducing better descriptions of such spatial effects as those observed in *Effects of Diffusivity* to non-spatial models. To this end, cellularization provides the ability to generate non-spatial models to describe various “compartments” of a biological system (*e.g.*, upper and lower respiratory tract compartments), where the non-spatial model well describes the underlying biocomplexity of a compartment according to its local specificity. We envision broad application of cellularization to do detailed spatiotemporal modeling of a particular biological module and then propagate information to, from and across coarser scales. Indeed, previous work has cast such a vision concerning the hierarchical organization and function of the liver (34). Cellularization provides a clear and consistent approach to develop such multiscale modeling frameworks of whole organisms with regards to the scale of the cell.

## Conclusion

In this work we developed a method for generating spatial, multicellular models of biological systems from non-spatial models, and *vice versa*, which we call cellularization. We demonstrate using our method by cellularizing non-spatial models of viral infection and host-pathogen interaction. Using these cellularized models, we quantitatively showed how spatial mechanisms implicitly represented in non-spatial models can exhibit significant effects on emergent dynamics when explicitly modeled. Variations in related non-spatial model parameters emerged from moderate cases of varying spatial mechanisms like rate of diffusive mass transport of virus, while extreme cases generated emergent dynamics inconsistently with those described by the non-spatial model. We describe the responsible mechanisms for extreme disagreement between homogeneous and cellular models, specifically concerning the limitations of describing discrete biological objects and processes using continuous descriptions.

## Acknowledgments

The authors acknowledge funding from National Institutes of Health grants U24 EB028887 and R01 GM122424 and National Science Foundation grant NSF 1720625. This research was supported in part by Lilly Endowment, Inc., through its support for the Indiana University Pervasive Technology Institute. The funders had no role in manuscript preparation or the decision to submit the work for publication.

